# Genetically perturbed myelin as a risk factor for neuroinflammation-driven axon degeneration

**DOI:** 10.1101/2023.07.17.549427

**Authors:** Janos Groh, Tassnim Abdelwahab, Yogita Kattimani, Michaela Hörner, Silke Loserth, Viktoria Gudi, Robert Adalbert, Fabian Imdahl, Antoine-Emmanuel Saliba, Michael Coleman, Martin Stangel, Mikael Simons, Rudolf Martini

## Abstract

Axon degeneration and functional decline in myelin diseases are often attributed to loss of myelin but their relation is not fully understood. Perturbed myelinating glia can instigate chronic neuroinflammation and contribute to demyelination and axonal damage. Here we study mice with distinct defects in the proteolipid protein 1 gene that develop axonal damage which is driven by cytotoxic T cells targeting myelinating oligodendrocytes. We show that persistent ensheathment with perturbed myelin poses a risk for axon degeneration, neuron loss, and behavioral decline. We demonstrate that CD8^+^ T cell-driven axonal damage is less likely to progress towards degeneration when axons are efficiently demyelinated by activated microglia. Mechanistically, we show that cytotoxic T cell effector molecules induce cytoskeletal alterations within myelinating glia and aberrant actomyosin constriction of axons at paranodal domains. Our study identifies detrimental axon-glia-immune interactions which promote neurodegeneration and possible therapeutic targets for disorders associated with myelin defects and neuroinflammation.

## Introduction

Myelination of axons by oligodendrocytes in the central nervous system (CNS) enables saltatory conduction, provides metabolic and trophic support, and modulates experience-dependent signal transmission^1, 2^. However, these evolutionary benefits are associated with an increased vulnerability of white matter to various pathogenic processes including disease- and aging-related immune reactions^3–8^. Dysfunctional interactions between neural and immune cells are increasingly recognized to initiate and perpetuate neuroinflammation, contributing to white matter pathology and neurological dysfunction^9–11^. Oligodendrocyte-lineage cells are immunocompetent glial cells and actively participate in these processes^8, 12–14^. Accordingly, chronic neuroinflammation is known to modify disorders associated with myelin defects and neurodegeneration, such as multiple sclerosis and hereditary leukodystrophies but also aging-related diseases like Alzheimer’s and Parkinson’s^15–18^. Axon degeneration in such disorders is most often proposed to be a consequence of chronic myelin and oligodendrocyte loss and the increased vulnerability of denuded axons to a toxic microenvironment^19–21^. However, the relationship between immune reactions, demyelination, axon degeneration, and clinical disease is unclear, and many observations indicate that loss of myelin itself does not correlate well with progressive neurodegeneration^21–24^.

We have previously demonstrated that gene defects perturbing the major CNS myelin proteolipid protein (PLP) - implicated in leukodystrophy and multiple sclerosis - result in neuroinflammation which amplifies neural damage and represents a target for treatment strategies^16^. In mice overexpressing normal or carrying mutant PLP (*PLPtg* and *PLPmut* mice, respectively) cytotoxic CD8^+^ T cells accumulate in the white matter and contribute to axon and myelin damage^25, 26^. After cognate T cell receptor (TCR) engagement within the CNS, these cells cause an impairment of axonal transport and a formation of axonal spheroids at juxtaparanodal domains of the nodes of Ranvier^27–29^. Moreover, innate immune reactions by microglia within the white matter contribute to neuroinflammation and myelin degradation in both disease models^30, 31^. Despite these commonalities, the exact mechanisms by which cytotoxic T cells drive damage of myelinated axons are unclear and several independent studies have indicated that neural-immune interactions might be associated with distinct disease outcomes in these mice. While *PLPtg* mice predominantly develop a demyelinating phenotype with periaxonal vacuole formation and some other axonopathic alterations of unknown clinical impact^25, 32^, *PLPmut* mice show chronic progressive neurodegeneration, clinical disease, and comparatively mild demyelination^26, 29, 33^. Therefore, these models of rare hereditary diseases caused by defects within the same myelin gene offer unique opportunities to study how axon degeneration, demyelination and behavioral deficits are related in chronic neuroinflammation, with important implications for more common disorders.

Here we directly compare the progression of clinical and histopathological characteristics between *PLPmut* and *PLPtg* mice. In contrast to the widely accepted model that demyelination leads to axon loss, we find an inverse relationship of axon degeneration and demyelination. We characterize neural-immune interactions in both models and identify shared and distinct glial reactions. Using experimental approaches to manipulate microglial myelin phagocytosis, we observe that persistent ensheathment with genetically perturbed myelin is a risk factor for neuroinflammation-driven axon degeneration and subsequent neuron loss. Along these lines, we uncover a novel mechanism of how myelinated axons are constricted at the paranodal domains as a reaction to the glia directed CD8^+^ T cell attack.

## Results

### Persisting ensheathment with perturbed myelin is associated with T cell-driven axon degeneration and neuron loss

Mice carrying mutant (*PLPmut*) or overexpressing normal (*PLPtg*) PLP have been reported to display similarities and differences in pathogenesis and disease outcome (Table 1). Due to these characteristics, we set out to address the following questions in the disease models: How are axon degeneration and demyelination related to disability and to each other? How are overlapping but distinct immune reactions related to axon degeneration and demyelination? How do glia directed immune reactions drive axonal damage and degeneration? To characterize the functional impact of the distinct perturbations affecting myelinating glia we used independent behavioral readout measures representing different neural circuits. While *PLPmut* mice (R137W, homozygous^26^) showed a significant decline in rotarod performance compared with non-transgenic wildtype (*Wt*) mice between 12 and 18 months of age, *PLPtg* mice (line 66, hemizygous^34^) retained normal motor coordination (Fig. 1a).

**Fig. 1.**
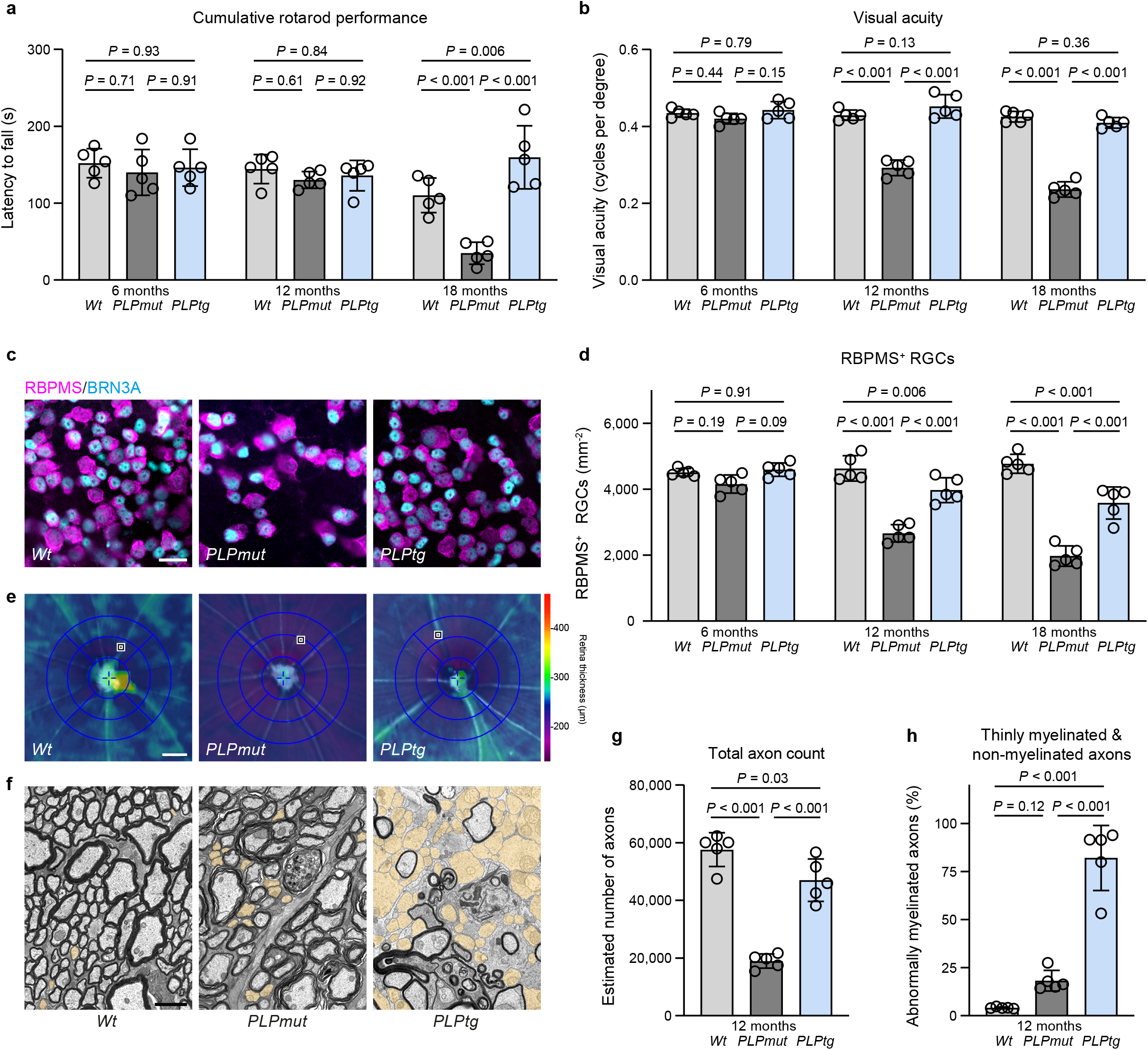
Neurodegeneration and myelin loss correlate inversely in mice with distinct myelin defects. a. Accelerating rotarod analysis of motor performance in *Wt*, *PLPmut*, and *PLPtg* mice (each circle represents the mean value of five consecutive runs of one mouse) at different ages. Motor performance is significantly impaired in *PLPmut* but not *PLPtg* mice at advanced disease stage (*n* = 5 mice per group, two-way ANOVA with Tukey’s multiple comparisons test, *F* (2, 36) = 14.88, *P* < 0.001). **b** Automated optokinetic response analysis of visual acuity (cycles per degree) shows a progressive decline of visual acuity in *PLPmut* but not *PLPtg* mice (each circle represents the mean value of one mouse, *n* = 5 mice per group, two-way ANOVA with Tukey’s multiple comparisons test, *F* (2, 36) = 195.0, *P* = 0.001). **c** Immunofluorescence detection and **d** quantification of RBPMS^+^BRN3A^+^ RGC in the retinae of *Wt*, *PLPmut*, and *PLPtg* mice. Neuron loss is much more pronounced in *PLPmut* than in *PLPtg* mice (*n* = 5 mice per group, two-way ANOVA with Tukey’s multiple comparisons test, *F* (2, 36) = 112.3, *P* < 0.001. Scale bar, 20 μm. **e** Representative interpolated thickness maps from retinal volume scans of 12-month- old *Wt*, *PLPmut*, and *PLPtg* mice. Scale bar corresponds to 50 μm of subtended retina. **f** Representative electron micrographs of optic nerve cross-sections from 12- month-old *Wt*, *PLPmut*, and *PLPtg* mice (thinly and non-myelinated axons are indicated in yellow pseudocolor). Scale bar, 2 μm. **g** Electron microscopy-based estimation of total axonal numbers in the optic nerves. Axon loss is much milder in *PLPtg* than in *PLPmut* mice (*n* = 5 mice per group, one-way ANOVA with Tukey’s multiple comparisons test, *F* (2, 12) = 63.67, *P* < 0.001). **h** Quantification of thinly myelinated (g-ratio ≥ 0.85) and non-myelinated axons in *Wt*, *PLPmut*, and *PLPtg* mice shows more prominent myelin loss in *PLPtg* mice (*n* = 5 mice per group, one- way ANOVA with Tukey’s multiple comparisons test, *F* (2, 12) = 81.47, *P* < 0.001). Data are presented as the mean ± s.d.

**Table 1.**
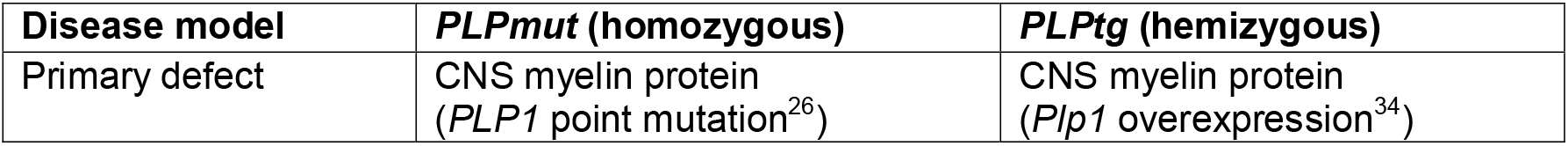
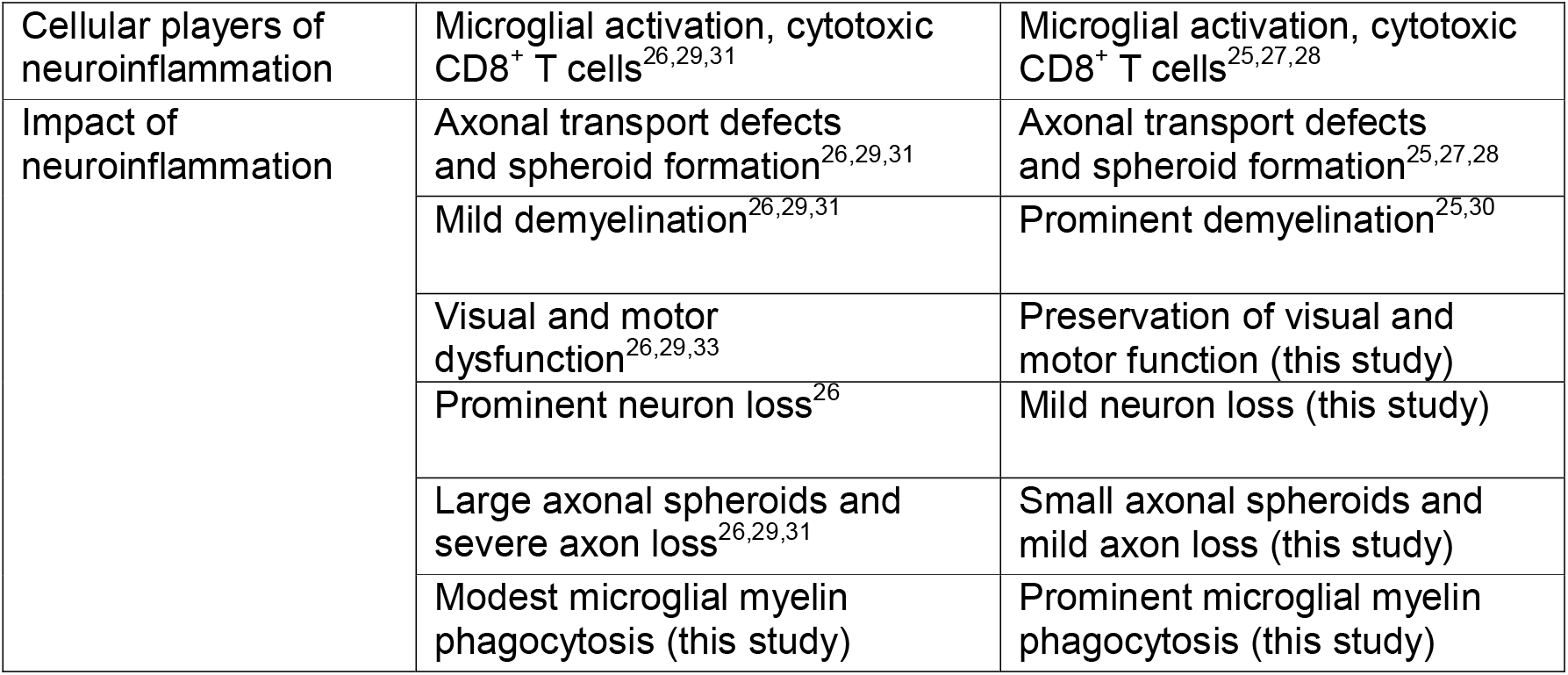
I Similarities and differences in pathogenesis and disease outcome of mice carrying mutant (*PLPmut*) compared with mice overexpressing normal (*PLPtg*) PLP.

Similarly, *PLPmut* but not *PLPtg* mice demonstrated a progressive decline in visual acuity that started even before 12 months of age (Fig. 1b). Structurally, this was accompanied by a much more pronounced loss of retinal ganglion cells in *PLPmut* compared with *PLPtg* mice (Fig. 1c,d). Optical coherence tomography (OCT) also demonstrated earlier and more prominent thinning of the innermost retinal layers in *PLPmut* mice (Fig. 1e, Supplementary Fig. 1a), arguing for more severe neuroaxonal degeneration. Electron microscopic quantification confirmed more axonopathic profiles (Supplementary Fig. 1b) and a severe loss of axons in optic nerves of *PLPmut* mice at 12 months of age, while *PLPtg* mice showed only a mild loss (Fig. 1f,g). In contrast, the frequency of thinly or non-myelinated axons was significantly higher in *PLPtg* compared with *PLPmut* mice (Fig. 1f,h). However, both disease models showed similar dynamics regarding the formation of axonal spheroids, which we labelled using antibodies against non-phosphorylated neurofilaments (SMI32, Supplementary Fig. 1c). Axonal spheroid formation was associated with an accumulation of CD8^+^ T cells in the white matter parenchyma of both models (Supplementary Fig. 1d-f), in line with their previously identified causal involvement in axonal damage^25, 26, 28, 29^. In summary, we detected an inverse relationship between neurodegeneration (associated with functional decline) and demyelination when comparing the two mouse models, despite an initially similar T cell-driven damage of myelinated axons.

We then compared the formation and fate of axonal spheroids in more detail. Electron microscopy and immunohistochemistry indicated that axonal spheroids in *PLPmut* mice were more often of larger size and still myelinated, while axonal spheroids appeared smaller and frequently non-myelinated in *PLPtg* mice (Fig. 2a,b). Detailed analyses at multiple ages revealed that axonal spheroids with small diameter (1.5 to 4 µm) form early in both models and are detectable at all investigated ages (Fig. 2c). However, axonal spheroids frequently progressed to larger sizes (> 4 µm diameter) in *PLPmut* mice, especially at advanced disease stages (12 and 18 months of age) while this was not the case in *PLPtg* mice. Additionally, we observed that axonal spheroids initially are still mostly myelinated in both models and become demyelinated with disease progression in *PLPtg* but rarely in *PLPmut* mice (Fig. 2d,e, Supplementary Fig. 1e). Thus, neuroinflammation-related axonal spheroids initially form similarly within perturbed myelin segments, but they only appear to progress in size and ultimately lead to axon degeneration when they remain myelinated. In line with these observations, CD8^+^ T cells in *PLPtg* mice preferentially associated with SMI32^+^ fibers that were still myelinated at an age when around 80% of optic nerve axons are demyelinated (Supplementary Fig. 2). These observations argue against the view that loss of myelin makes axons susceptible to secondary immune-mediated degeneration and instead imply that efficient demyelination may allow resilience of axons at reversible stages of damage.

**Fig. 2.**
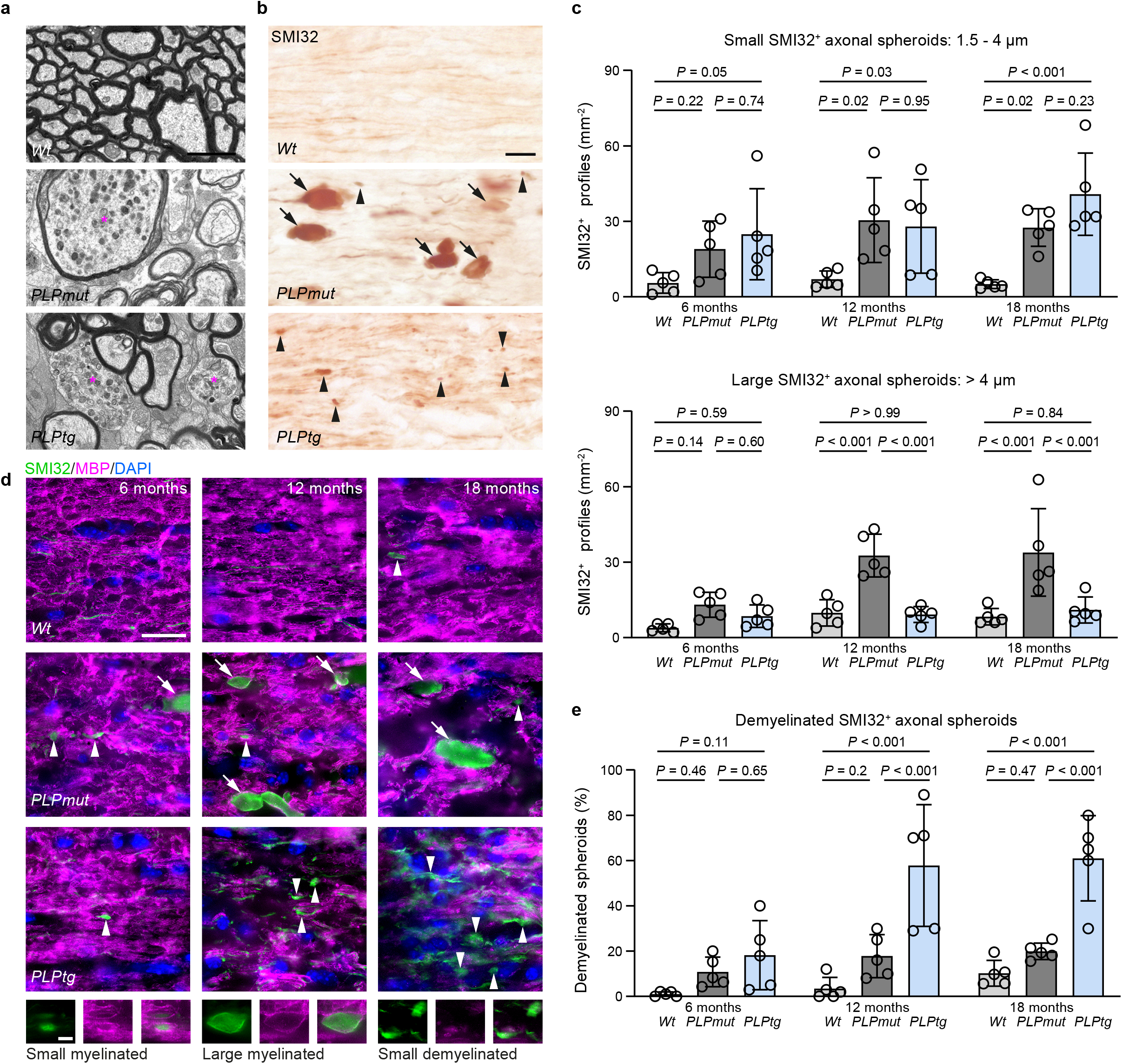
Axonal spheroids show a similar initial formation but a different progression in distinct myelin mutants. a. Representative electron micrographs of optic nerve cross-sections from 12-month-old *Wt*, *PLPmut*, and *PLPtg* mice demonstrate differences in size and myelination state of axonal spheroids (asterisks). Scale bar, 2 μm. **b** Immunohistochemical visualization of SMI32^+^ axonal spheroids in the optic nerves of 12-month-old *Wt*, *PLPmut*, and *PLPtg* mice. Arrowheads indicate small axonal spheroids (diameter 1.5 - 4 µm) and arrows indicate large axonal spheroids (diameter > 4 µm). Scale bar, 20 μm. **c** Quantification of small (top) and large (bottom) axonal spheroids at different ages (each circle represents the mean value of one mouse). Small spheroids are similarly frequent in *PLPmut* and *PLPtg* mice but become large with disease progression only in *PLPmut* mice (*n* = 5 mice per group, two-way ANOVA with Tukey’s multiple comparisons test, Small: *F* (2, 36) = 16.71, *P* < 0.001, Large: *F* (2, 36) = 30.04, *P* < 0.001). **d** Immunofluorescence detection of SMI32^+^ axonal spheroids (small: arrowheads, large: arrows) and MBP in the optic nerves of *Wt*, *PLPmut*, and *PLPtg* mice at different ages. Scale bar, 20 μm. Bottom images show examples of spheroids of different size and myelination state with split channels at higher magnification. Scale bar, 5 µm. **e** Quantification of the myelination state of SMI32^+^ axonal spheroids demonstrates their progressive demyelination in *PLPtg* but not *PLPmut* mice (*n* = 5 mice per group, two-way ANOVA with Tukey’s multiple comparisons test, *F* (2, 36) = 39.45, *P* < 0.001). Data are presented as the mean ± s.d.

### Distinct neural-immune interactions determine the demyelination of axons ensheathed with perturbed myelin

To better characterize the cellular interactions that determine these different outcomes regarding the persistence of perturbed myelin, we investigated transcriptional changes within myelinating glia and CNS-resident phagocytes. We therefore used single-cell RNA sequencing (scRNA-seq) of CD45^-^O1^+^ mature oligodendrocytes and CD45^low^Siglec-H^+^ microglia sorted from the brains of adult (10- month-old) *Wt*, *PLPmut*, and *PLPtg* mice (gating strategy presented in Supplementary Fig. 3a). After quality control and filtering (Supplementary Fig. 3b), unsupervised clustering of the combined datasets identified nine different microglia clusters and a smaller oligodendrocyte (ODC) cluster (Fig. 3a, Supplementary Table 1). Within ODC, we detected shared and unique disease-related transcriptional alterations in the distinct models. Both in *PLPmut* and *PLPtg* mice several transcripts indicating altered proteostasis and cell stress (e.g., *Uba52*, *Cd63*, *Rps5*), and inflammatory signaling (e.g., *Il33*, *Ccl4*, *Lyz2*) were differentially expressed, likely as a direct consequence of the distinct myelin gene defects (Supplementary Fig. 3c,d, Supplementary Table 1). However, while ODC in *PLPmut* mice upregulated several transcripts related to protection against cell death and phagocytosis (e.g., *Cryab*, *Xiap*, *Cd47*), ODC in *PLPtg* mice expressed high levels of phagocytosis-promoting transcripts (e.g., *C1qa*, *C1qb*, *Apoe*).

**Fig. 3.**
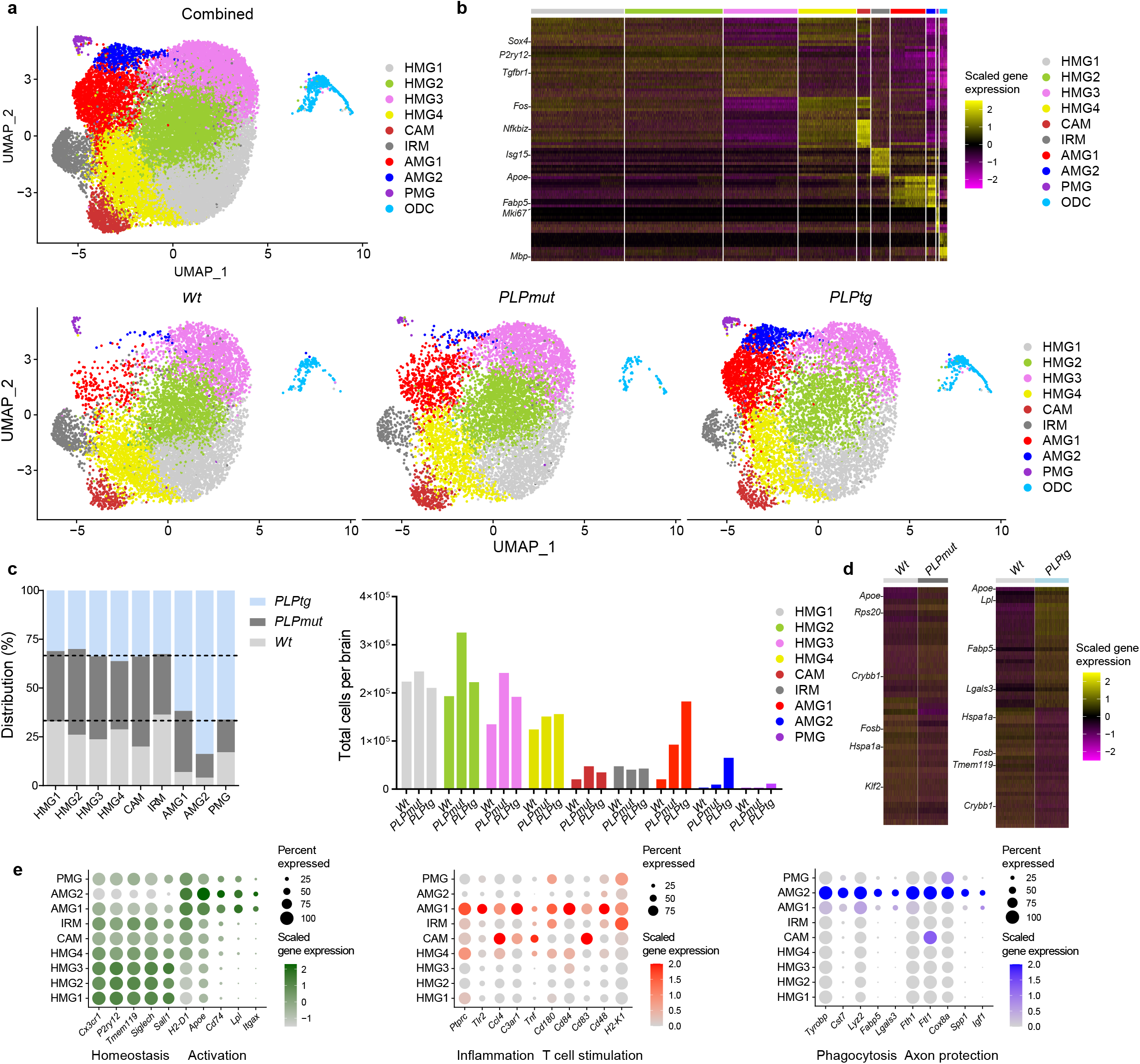
scRNA-seq reveals heterogeneous neural-immune interactions in mice with distinct myelin defects. a. UMAP visualization of CD45^-^O1^+^ mature oligodendrocytes and CD45^low^Siglec-H^+^ microglia freshly sorted from adult (10- month-old) *Wt*, *PLPmut*, and *PLPtg* (*n* = 3 mice per group) mouse brains and analyzed by scRNA-seq. Combined (top, 26,308 cells) and separate (bottom) visualization of cells from *Wt* (9,484 cells), *PLPmut* (8,405 cells), and *PLPtg* (8,419 cells) brains are displayed. **b** Heatmap of top 10 cluster-specific genes. The color scale is based on a z-score distribution from -2 (purple) to 2 (yellow). **c** Contribution of the samples to each microglia cluster is displayed in percent (left) and absolute numbers extrapolated to total cells per brain (right). AMG1 is enriched in both myelin mutants and AMG2 mostly in *PLPtg* mice. **d** Heatmaps of top 30 differentially expressed genes comparing microglia isolated from *Wt* and *PLPmut* (left) or *Wt* and *PLPtg* (right) brains across all clusters as identified in panel a. **e** Dot plot expression visualization of selected genes implicated in microglial homeostasis and activation (left), inflammation and T cell stimulation (middle), or phagocytosis and axon protection (right) for microglia clusters as annotated in panel a. The color scales are based on z-score distributions from -1 (lightgrey) to 2 (green) or 0 (lightgrey) to 2 (red, blue). HMG, homeostatic microglia; CAM, capillary-associated microglia; IRM, interferon-responsive microglia; AMG, activated microglia; PMG, proliferating microglia; ODC, oligodendrocytes. Complete lists of cluster-specific marker and differentially expressed genes can be found in Supplementary Table 1.

We next focused on the consequences of these alterations for microglial diversity and function. Four of the microglia clusters resembled homeostatic microglia (HMG1- 4; Fig. 3b)^35^. Moreover, we identified one cluster with a signature enriched in immediate early genes and *Icam1* and one cluster with a prominent interferon response signature, reminiscent of previously described capillary-associated microglia (CAM, see below) and interferon-responsive microglia (IRM), respectively^36, 37^. We also detected the presence of two groups of activated microglia (AMG1-2) with partially overlapping transcriptional signatures and a small population of proliferating microglia (PMG). Both AMG clusters showed transcriptional similarities to previously described disease-, aging-, or neuroinflammation-related microglia states such as DAM^38^, MGnD^39^, WAM^40^, microglia in EAE^41^, after systemic LPS challenge^42^, and to a smaller extend DIM^43^ (which showed more transcriptional similarities to CAM; Supplementary Fig. 4a, Supplementary Table 1). Moreover, AMG and PMG were strongly enriched in the disease models, with AMG2 and PMG being more unique to *PLPtg* mice (Fig. 3c). Global myelin disease-related transcriptional changes in microglia were primarily based on increased numbers of cells representing AMG and reflected the prominent contribution of AMG2 in *PLPtg* mice (Fig. 3d). Focusing on the activated microglia states, we found that both shared (to different degrees) the typical downregulation of markers indicating homeostatic function (e.g., *Cx3cr1*, *P2ry12*, *Tmem119*, *Siglech*, *Sall1*) and the upregulation of activation markers (e.g., *H2-D1*, *Apoe*, *Cd74*, *Lpl*, *Itgax*) reminiscent of the DAM program^44^ (Fig. 3e, Supplementary Fig. 4b). However, when looking for subset-enriched signatures (marker genes that are also significantly upregulated in one vs the other AMG cluster), we observed that AMG1 expressed higher levels of genes associated with pro-inflammatory signaling (e.g., *Ptprc*, *Tlr2*, *Ccl4*, *C3ar1*, *Tnf*) and T cell stimulation (e.g., *Cd180*, *Cd84*, *Cd83*, *Cd48*, *H2-K1*), whereas AMG2 showed enrichment of genes related to phagocytosis (e.g., *Tyrobp*, *Cst7*, *Lyz2*, *Fabp5*, *Lgals3*) and axon protection (e.g., *Fth1*, *Ftl1*, *Cox8a*, *Spp1*, *Igf1*). GO analysis reflected these differences between AMG states (Supplementary Fig. 4c). Flow cytometry of CD45^low^Siglec-H^+^ microglia from brains of 12-month-old *Wt*, *PLPmut*, and *PLPtg* mice using identified markers confirmed the presence of distinct populations and increased frequency of the AMG subsets in both disease models (Supplementary Fig. 5). Using CD11c (encoded by *Itgax* and specific to both AMG clusters) and PD-1 (encoded by *Pdcd1* and specific to AMG1) as discriminating markers, we further confirmed the stronger accumulation of AMG2 in *PLPtg* mice.

Immunohistochemistry revealed that AMG (CD11c^+^) specifically arise in the white matter of mice with myelin perturbation (Supplementary Fig. 6a,b). Moreover, CAM (ICAM1^+^) were localized in proximity to capillaries in adult *Wt* mice (Supplementary Fig. 6c). Interferon-responsive microglia and astrocytes were previously localized to white matter regions in proximity to the ventricles^8, 45, 46^. Enzymatic dissociation artifacts^47^ could not fully explain the detection of the distinct HMG subsets, since c-Jun expression levels (part of the *ex vivo* activation signature and differentially expressed between HMG subsets) varied between individual microglia in adult *Wt* mice (Supplementary Fig. 6d,e). In addition, MHC-I expression (differing between HMG1-2 and HMG3-4) was higher in *Wt* white matter compared with gray matter microglia (Supplementary Fig. 6d,f). Electron microscopy and immunohistochemistry showed that myelin-phagocytosing amoeboid microglia with an accumulation of lysosomal storage material were frequent in *PLPtg* but not *PLPmut* mice (Fig. 4a,b). Immunohistochemical quantification confirmed that the total number of microglia (CD11b^+^) as well as AMG (CD11c^+^) was increased in optic nerves of both disease models. However, while AMG1 (P2RY12^+^CD11c^+^) was enriched in both models, AMG2 (P2RY12^-^ CD11c^+^) was specific to *PLPtg* mice (Fig. 4c-e). In line with our scRNA-seq data, galectin 3 (encoded by *Lgals3*), associated with phagocytosis^48^, was detectable at higher levels in AMG2 (P2RY12^-^CD11c^+^) of *PLPtg* mice than in AMG1 (P2RY12^+^CD11c^+^) of *PLPmut* or HMG (P2RY12^+^CD11c^-^) of *Wt* mice. In summary, distinct myelin defects within oligodendrocytes of *PLPmut* and *PLPtg* mice appear to drive similar pro-inflammatory microglial activation (represented by AMG1). However, they result in varying responses regarding the transition into a myelin-phagocytosing microglia state (represented by AMG2). These reactions might depend on intrinsic differences related to the myelin gene defects or represent distinct responses to the glia directed CD8^+^ T cell attack.

**Fig. 4.**
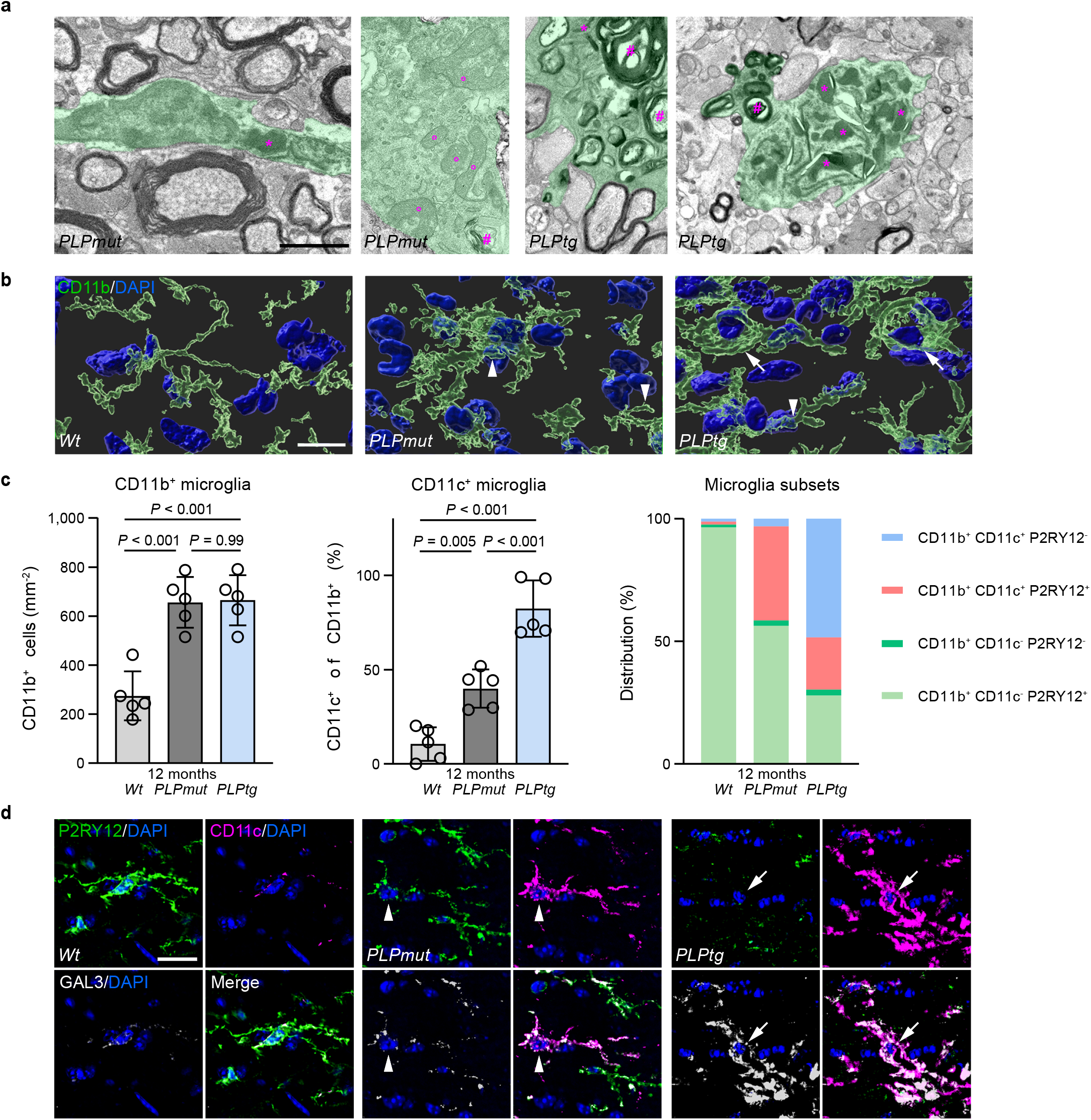
Myelin phagocytosis by activated microglia mediates demyelination in mice with distinct myelin defects. a. Representative electron micrographs of microglial cells (green pseudocolor) in optic nerve cross-sections from 12-month-old *PLPmut*, and *PLPtg* mice demonstrate differences in morphology, mitochondrial content (circles), and intracellular accumulation of myelin fragments (hashtags) and lysosomal storage material (asterisks). Scale bar, 2 μm. **b** Immunofluorescence detection and IMARIS Z-stack surface rendering of CD11b^+^ microglia in optic nerves of *Wt*, *PLPmut*, and *PLPtg* mice. Arrowheads indicate “bushy” microglia with thick processes and arrows indicate “amoeboid” microglia with short processes. Scale bar, 10 μm. **c** Quantification of CD11b^+^ microglia (left), CD11c^+^ microglia (% of CD11b, middle), and distribution of CD11c/P2RY12 reactivity on microglia (% of CD11b, right) in the optic nerves of 12-month-old *Wt*, *PLPmut*, and *PLPtg* mice (each circle represents the mean value of one mouse) as shown in Supplementary Fig. 6a,b. Ramified CD11c^+^ P2RY12^+^ microglia (representing AMG1) accumulate in both myelin mutants while amoeboid CD11c^+^ P2RY12^-^ microglia (representing AMG2) arise in *PLPtg* mice (*n* = 5 mice per group, one-way ANOVA with Tukey’s multiple comparisons test, Left: *F* (2, 12) = 23.88, *P* < 0.001, Middle: *F* (2, 12) = 48.00, *P* < 0.001). **d** Representative immunofluorescence detection of P2RY12, CD11c, and GAL3 in the optic nerves of 12-month-old *Wt*, *PLPmut*, and *PLPtg* mice. CD11c^+^ P2RY12^-^ (AMG2, arrow) microglia in *PLPtg* mice show higher expression of GAL3 than CD11c^+^ P2RY12^-^ (AMG1, arrowhead) microglia in *PLPmut* or CD11c^-^ P2RY12^+^ (HMG) microglia in *Wt* mice. Scale bar, 10 µm. Data are presented as the mean ± s.d.

### Phagocytosis of perturbed myelin by activated microglia counteracts T cell-driven axon degeneration

To further test the hypothesis that persisting ensheathment with perturbed myelin makes axons susceptible to neuroinflammation-related damage and degeneration in the distinct models, we used two reciprocal experimental approaches aiming to modulate demyelination. First, we fed *PLPmut* mice with the copper chelator cuprizone to promote toxin-based demyelination known to be driven by CSF-1- activated microglia^49, 50^. Second, we pharmacologically depleted CNS-resident myeloid cells with the CSF-1R inhibitor PLX5622 to attenuate microglia-driven demyelination in *PLPtg* mice^51^. These approaches were selected to foster or mitigate the predicted impact of microglial myelin phagocytosis based on their observed activation states and our previous work in the respective models^30, 31^. Electron microscopy-based quantification revealed a significant increase in thinly and non-myelinated axons in optic nerves of *PLPmut* mice after 6 weeks of cuprizone treatment (Fig. 5a). This was associated with fewer axons showing pathological alterations in form of spheroid formation and degenerative signs (Fig. 5b) and resulted in an attenuation of axon loss (Supplementary Fig. 7a). Moreover, cuprizone treatment increased the frequency of *Csf1*^+^*Mbp*^+^ oligodendrocytes and amoeboid microglia with lysosomal storage material in *PLPmut* mice (Supplementary Fig. 7b,c). On the other hand, 6 months of continuous PLX5622 treatment resulted in a reduced frequency of thinly- and non-myelinated axons and an increased frequency of axonal spheroids and degenerating axons in *PLPtg* but not *Wt* mice (Fig. 5c,d). This led to an aggravated loss of axons and retinal ganglion cells (Supplementary Fig. 7d,e). To study the consequences of even faster removal of perturbed myelin on axon integrity, we also investigated homozygous *PLPtg* mice. In contrast to hemizygous mice, these showed almost complete absence of myelin in the optic nerves at 4 months of age without detectable axon loss (Supplementary Fig. 7f,g). Corroborating our observations regarding myelin phagocytosis, cuprizone treatment led to a significant increase in the number of galectin 3^+^ microglia^48^ (encoded by *Lgals3* - enriched within AMG2) in *PLPmut* mice, while PLX5622 decreased their density in *PLPtg* mice (Fig. 5c). These observations indicate that fostering microglia-mediated demyelination attenuates axonal degeneration and that impairing demyelination aggravates axonal degeneration when perturbed myelin is targeted by adaptive immunity.

**Fig. 5.**
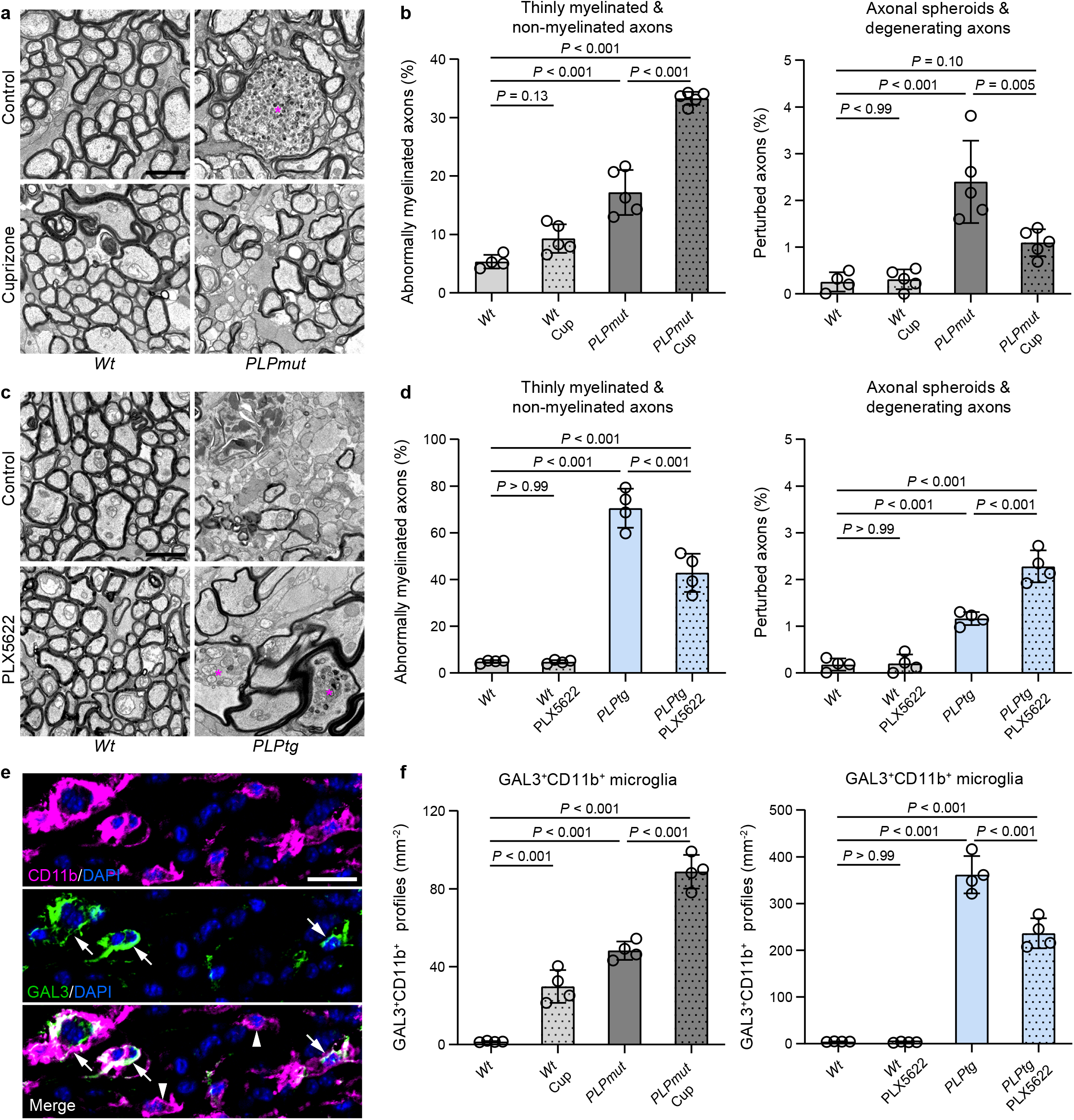
Modulating myelin phagocytosis inversely affects axonal damage in distinct myelin mutants. a. Representative electron micrographs of optic nerve cross-sections from control (top) or cuprizone treated (bottom) *Wt* (left) and *PLPmut* (right) mice. The asterisk indicates an axonal spheroid. Scale bar, 2 μm. **b** Electron microscopy-based quantification of thinly myelinated (g-ratio ≥ 0.85) and non- myelinated axons (left) or axonal spheroids and degenerating axons (right) in *Wt* and *PLPmut* mice (*n* = 5 mice per group) after control or cuprizone (Cup) diet (each circle represents the mean value of one mouse). Cuprizone significantly fosters myelin loss and attenuates axonal damage in *PLPmut* but not *Wt* mice (*n* = 4-5 mice per group, one-way ANOVA with Tukey’s multiple comparisons test, Left: *F* (3, 15) = 117.5, *P* < 0.001, Right: *F* (3, 15) = 19.13, *P* < 0.001). **c** Representative electron micrographs of optic nerve cross-sections from control (top) or PLX5622 treated (bottom) *Wt* (left) and *PLPtg* (right) mice. The asterisks indicate axonal spheroids. Scale bar, 2 μm. **d** Electron microscopy-based quantification of thinly myelinated (g-ratio ≥ 0.85) and non-myelinated axons (left) or axonal spheroids and degenerating axons (right) in *Wt* and *PLPtg* mice (*n* = 4 mice per group) after control or PLX5622 diet. PLX5622 significantly attenuates myelin loss and fosters axonal damage in *PLPtg* but not *Wt* mice (*n* = 4 mice per group, one-way ANOVA with Tukey’s multiple comparisons test, Left: *F* (3, 12) = 118.8, *P* < 0.001, Right: *F* (3, 12) = 82.59, *P* < 0.001). **e** Representative immunofluorescence detection of GAL3 reactivity of CD11b^+^ microglia in optic nerves. Arrowheads indicate GAL3^-^ microglia and arrows indicate GAL3^+^ microglia. Scale bar, 20 μm. **f** Quantification of GAL3^+^CD11b^+^ microglia in the optic nerves of *Wt* and *PLPmut* mice after control or cuprizone (Cup) diet (left) and *Wt* and *PLPtg* mice after control or PLX5622 diet (right). GAL3^+^ microglia (representing AMG2) accumulate in *PLPmut* mice after cuprizone diet and are reduced in number after PLX5622 diet in *PLPtg* mice (*n* = 4 mice per group, one-way ANOVA with Tukey’s multiple comparisons test, Left: *F* (3, 12) = 127.4, *P* < 0.001, Right: *F* (3, 12) = 194.9, *P* < 0.001). Data are presented as the mean ± s.d.

### Focal damage of axons ensheathed by perturbed myelin occurs in proximity to constricted paranodal regions

Next, we addressed the underlying mechanisms responsible for the formation of spheroids as early signs of focal damage by CD8^+^ T cells to axons ensheathed by perturbed myelin. Upon close examination of longitudinal optic nerve sections by electron microscopy, we confirmed the previously described preferential localization of spheroids at juxtaparanodal axon domains^26, 28^ and frequently observed an accumulation of disintegrating mitochondria and other organelles (an early sign of axonal damage^52, 53^). Strikingly, at the corresponding paranodal domains, axons appeared to be of smaller diameter than usual (Fig. 6a). In *PLPmut* mice, progressive swelling, accumulation of dense bodies within myelinated spheroids, and fragmentation appeared to correlate with decreasing paranodal diameters. In *PLPtg* but not *PLPmut* mice, we frequently detected phagocytosing microglia stripping early-stage axonal spheroids of myelin. Abnormally small paranodal axon diameters were also detectable in cross sections of *PLPmut* optic nerves, excluding misinterpretation related to oblique fiber orientation in longitudinal sections (Fig. 6b). This previously unknown narrowing of the paranodal axon diameter reminiscent of a constriction process could explain the described impairment of axonal transport and consequent juxtaparanodal accumulation of organelles in the disease models^26, 28^. We therefore measured the minimum axonal diameters of paranodal domains with adjacent spheroid formation categorized into different putative stages of progression (Fig. 6c). Normal appearing fibers in *Wt*, *PLPmut*, and *PLPtg* mice displayed larger paranodal axon diameters than those at stages 1 (accumulation of organelles) and 2 (swelling). An increased frequency especially of stage 2 profiles resulted in a significant reduction of the mean paranodal axon diameter in *PLPmut* compared with *Wt* and *PLPtg* mice. Moreover, in longitudinal sections, oligodendroglial cytoplasmic loops displayed decreased circularity going along with progressive spheroid formation (Supplementary Fig. 8a, Fig. 6d). This suggests that abnormal myelinating process extension may contribute to constriction of paranodal axon diameters. Focusing on *PLPmut* mice, we further studied the putative role of paranodal constriction in axon degeneration and tested its dependency on functional adaptive immunity. *Rag1*-deficient *PLPmut* mice (lacking mature adaptive immune cells) showed less decreased paranodal axon diameters (Supplementary Fig. 8b, Fig. 6e). We confirmed this immune-dependent overall decrease in paranodal axon diameters by measuring CASPR^+^ domains using immunofluorescence (Supplementary Fig. 8c). Moreover, in line with previous observations^26, 29^, absence of functional lymphocytes decreased the age-dependent formation of small and large axonal spheroids and axon loss in *PLPmut* mice (Fig. 6f,g). Thus, myelinated fibers under adaptive immune attack seem to constrict their diameters at paranodal domains and thereby hamper axonal transport which can promote axon degeneration^54^.

**Fig. 6.**
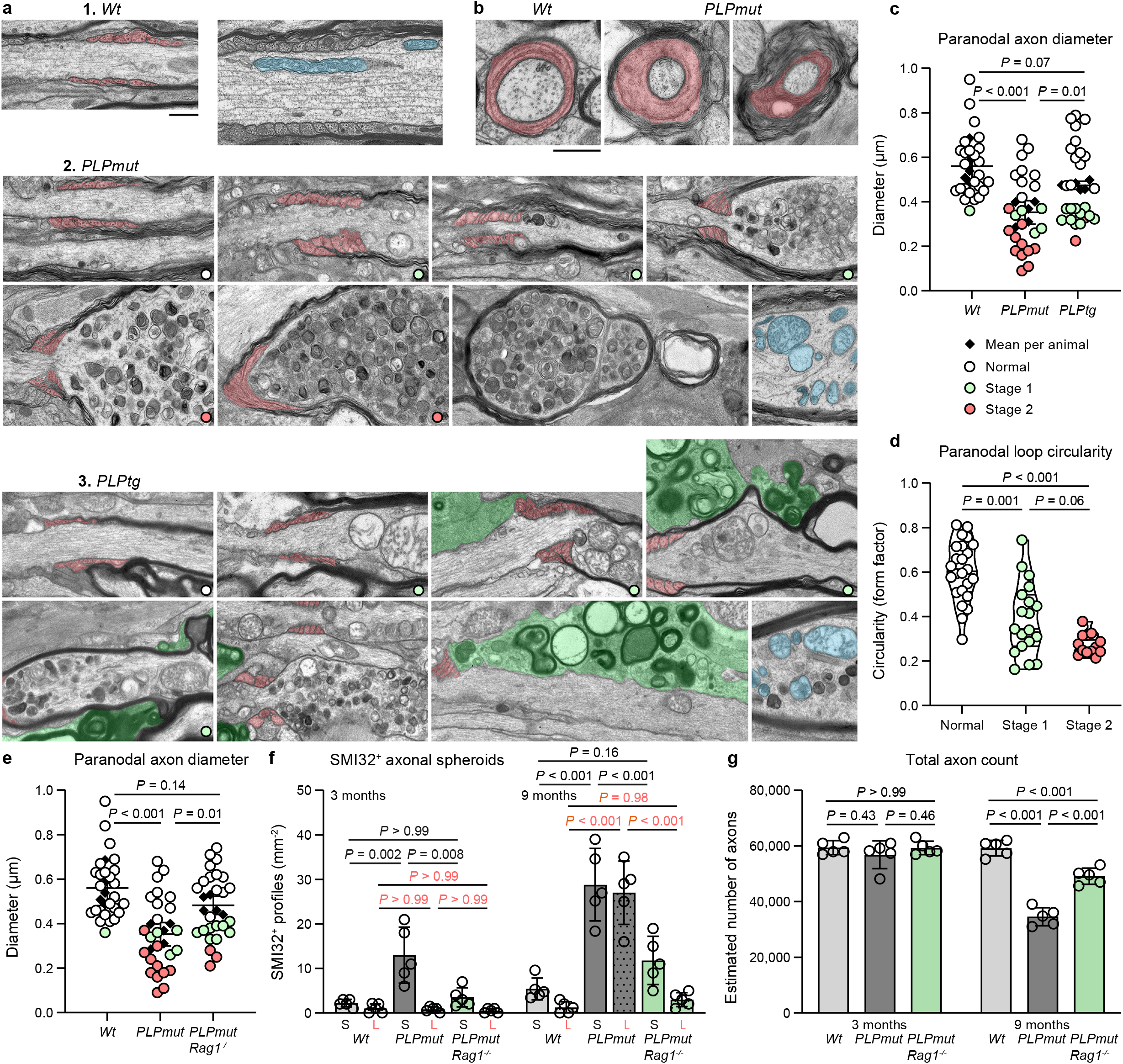
T cell-driven axonal spheroid formation is initiated in proximity to constricted paranodal domains. a. Representative electron micrographs of optic nerve longitudinal sections from 3- to 9-month-old *Wt* (top), *PLPmut* (middle), and *PLPtg* mice displaying putative subsequent stages of axonal spheroid formation. Progressive constriction of axon diameters by oligodendrocytic paranodal loops (light red pseudocolor) correlates with progressive accumulation of disintegrating mitochondria (exemplarily shown in light blue pseudocolor), dense bodies, juxtaparanodal swelling, and fragmentation in *PLPmut* mice. Early-stage axonal spheroids in *PLPtg* mice are often associated with myelin phagocytosing microglia (green pseudocolor) in *PLPtg* mice. Scale bar, 0.5 μm. **b** Fibers with constricted paranodal domains are also detectable in cross-sections of optic nerves from *PLPmut* mice. Scale bar, 0.5 µm **c** Electron microscopy-based quantification of minimum paranodal axon diameters in 9-month-old *Wt*, *PLPmut*, and *PLPtg* mice (each circle represents the value of one paranodal region and each diamond represents the mean value of one mouse, colors indicate stages of spheroid formation). Mean diameter is significantly reduced in *PLPmut* mice (*n* = 5 mice and 25 paranodes per group, one-way ANOVA with Tukey’s multiple comparisons test, *F* (2, 12) = 17.5, *P* < 0.001. **d** Analysis of the form factor of paranodal loops reveals decreased circularity along with progressive stages of spheroid formation (each circle represents the mean value of one paranodal region, *n* = 5 mice and 56 paranodes: 24 normal, 19 stage 1, 13 stage 2, one-way ANOVA with Tukey’s multiple comparisons test, *F* (2, 53) = 30.54, *P* < 0.001). **e** Electron microscopy- based quantification of minimum paranodal axon diameters in 9-month-old *Wt*, *PLPmut* (as shown in panel c), and *PLPmut*/*Rag1*^-/-^ mice. Reduction of the mean diameter in *PLPmut* mice is attenuated by *Rag1* deficiency (*n* = 5 mice and 25 paranodes per group, one-way ANOVA with Tukey’s multiple comparisons test, *F* (2, 12) = 15.55, *P* < 0.001). **f** Quantification of small (S) and large (L) SMI32^+^ axonal spheroids in 9-month-old *Wt*, *PLPmut*, and *PLPmut*/*Rag1*^-/-^ mice at different ages (each circle represents the mean value of one mouse). Small and large spheroids are significantly less frequent in *PLPmut*/*Rag1*^-/-^ compared with *PLPmut* mice (*n* = 5 mice per group, two-way ANOVA with Tukey’s multiple comparisons test, *F* (5, 48) = 36.19, *P* < 0.001). **g** Electron microscopy-based estimation of total axonal numbers in the optic nerves. Axon loss is much milder in *PLPmut*/*Rag1*^-/-^ than *PLPmut* mice (*n* = 5 mice per group, two-way ANOVA with Tukey’s multiple comparisons test, *F* (2, 24) = 44.33, *P* < 0.001). Data are presented as the mean ± s.d.

### Cytotoxic effector molecules induce cytoskeletal plasticity in myelinating oligodendrocytes

After peripheral nerve injury, Schwann cells form constricting actomyosin spheres to accelerate axon fragmentation in a Pak1-dependent and cytokinesis-like manner^55–57^. Moreover, oligodendrocytes show increased cytoskeletal dynamics after injury and some of the molecules important for myelinating process extension and ensheathment of axons are direct targets for cleavage by the T cell effector protease granzyme B (GZMB)^58–60^. We have previously demonstrated that immune-mediated axonal damage in *PLPmut* and *PLPtg* mice depends on GZMB^27, 29^ and therefore hypothesized that the focal cytotoxic attack on myelinating oligodendrocyte processes could induce cytoskeletal alterations within non-compacted myelin domains. Supporting this hypothesis, our scRNA-seq analysis of mature ODC from myelin mutants indicated increased expression of several factors related to membrane trafficking and plasma membrane repair (e.g., *Chmp2a*, *Vps4a*, *Syt11*, known to counteract T cell cytotoxicity^61^), regulation of actomyosin structure organization (e.g., *Rhoa*, *Cdc42*, *Pfn1*), and actin filament based process (e.g., *Pak1*, *Wasf2*, *Tpm3*) particularly in *PLPmut* mice (Supplementary Fig. 9a). Labelling of F-actin with phalloidin demonstrated increased levels at the outer aspects of myelin segments around cleaved caspase 3^+^ axonal spheroids in optic nerves of *PLPmut* mice, suggesting pro-degenerative signaling in axons likely constricted by oligodendrocytes (Supplementary Fig. 9b,c). Moreover, immunohistochemistry indicated increased diphosphorylated non-muscle myosin light chain (pMLC) levels in cell processes ensheathing SMI32^+^ axonal spheroids (Supplementary Fig. 9d), arguing for localized contractile myosin II activity within non-compact myelin domains. Like decreased paranodal diameter, increased pMLC levels in optic nerves of *PLPmut* mice were largely dependent on functional adaptive immunity (Supplementary Fig. 9e, Fig. 7a).

**Fig. 7.**
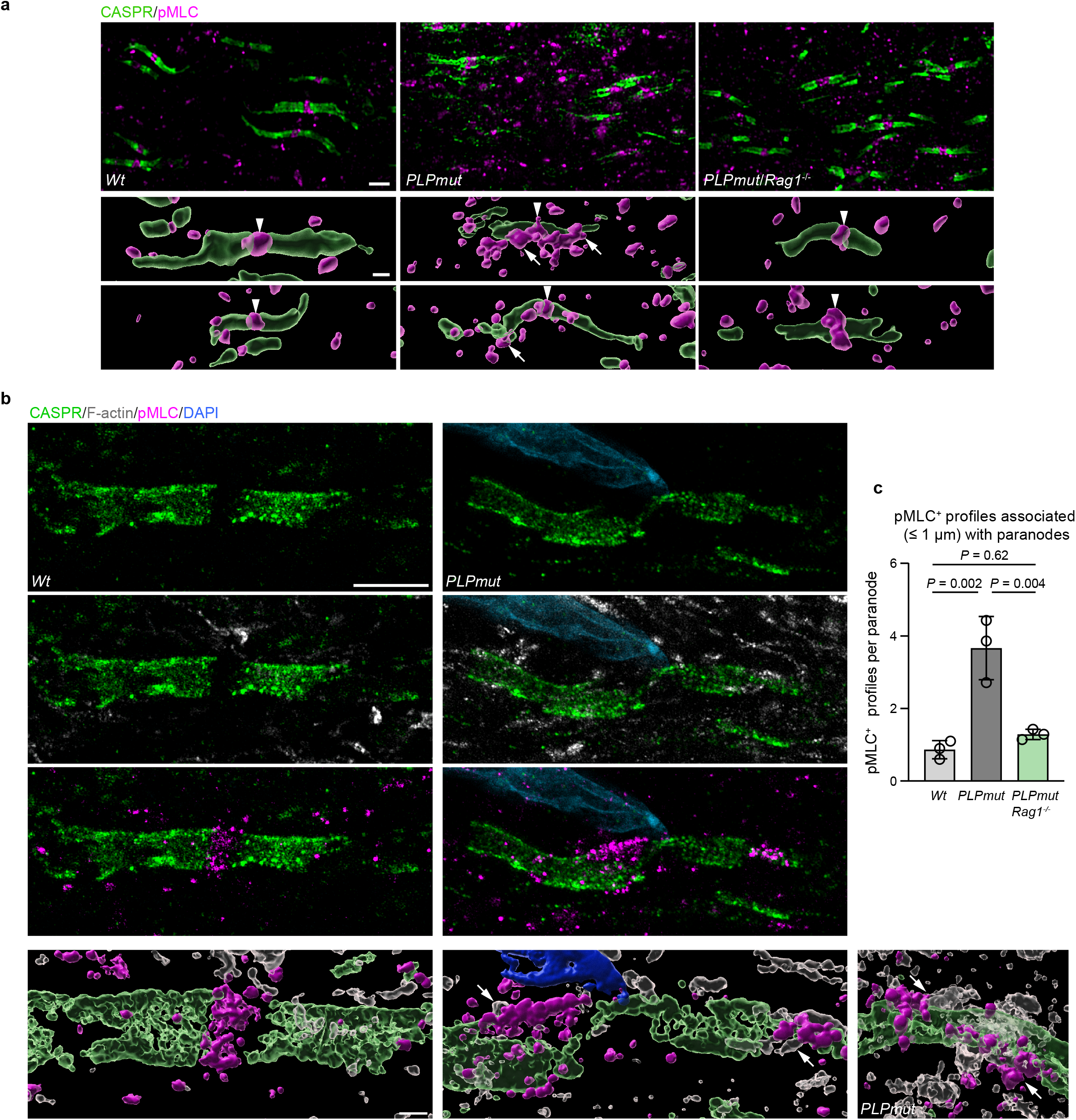
Constricted paranodal domains in myelin mutants are associated with increased actomyosin activity. a. Representative immunofluorescence detection and IMARIS Z-stack surface rendering of CASPR and pMLC in the optic nerves of 12-month-old *Wt*, *PLPmut*, and *PLPmut*/*Rag1*^-/-^ mice. pMLC is localized at normal appearing nodes of Ranvier (arrowheads). In *PLPmut* mice, paranodal domains with irregularly small diameter are surrounded by additional pMLC aggregates (arrows). Scale bar, 2 µm (top), 1 µm (bottom). **b** Super-resolution fluorescence detection and IMARIS Z-stack surface rendering of CASPR, F-actin, and pMLC in the optic nerves of 12-month-old *Wt* and *PLPmut* mice. Compact assemblies of F-actin and pMLC enwrap constricted paranodal domains in *PLPmut* mice (arrows). Note the close proximity of a cell nucleus to the constricted paranodal domain. Scale bars: 10 µm expanded, 2.5 µm unexpanded (top), 2 µm expanded, 500 nm unexpanded (bottom). **c** Quantification of pMLC^+^ profiles in close association (≤ 1 µm distance) with individual CASPR^+^ paranodes in optic nerves of *Wt*, *PLPmut*, and *PLPmut*/*Rag1*^-/-^ mice (each circle represents the mean value of one mouse). Paranodes in *PLPmut* mice are associated with a higher number of pMLC^+^ assemblies which is attenuated by *Rag1* deficiency (*n* = 25 paranodes per mouse and 3 mice per group, One-way ANOVA with Tukey’s multiple comparisons test, *F* (2, 6) = 24.27, *P* = 0.001). Data are presented as the mean ± s.d.

Since pMLC has been implicated in the maintenance of the membrane periodic skeleton of axons *in vitro*^62^, we localized the pMLC signal in more detail. Corroborating previous work^63^, pMLC was mostly confined to the nodal region flanked by CASPR^+^ paranodal domains in optic nerves of *Wt* mice (Fig. 7a). In contrast, CASPR^+^ paranodes in *PLPmut* but not *PLPmut*/*Rag1*^-/-^ mice were frequently surrounded by additional spots of pMLC reactivity, especially when showing focal signs of constriction (Fig. 7a,b). To confirm this, we used multiplexed super-resolution fluorescence microscopy by combining 4x physical expansion^64^ of optic nerve sections with confocal microscopy and an Airyscan 2 detector. This revealed focal narrowing of the paranodal domains in *PLPmut* mice which were frequently closely enwrapped by compact assemblies of F-actin and pMLC (Fig. 7b,c). Quantification of pMLC^+^ assemblies in close association (≤ 1 µm distance) with individual CASPR^+^ paranodes confirmed an immune-dependent accumulation in *PLPmut* mice (Fig. 7c). Since CASPR^+^ axonal domains are in direct contact with glial paranodal loops, this suggests local contractile actomyosin activity in non-compact domains of myelinating oligodendrocytes.

To further investigate the relationship between cytotoxic molecules and cytoskeletal dynamics we isolated OPCs from P7 mouse brains and differentiated them for seven days into myelinating oligodendrocytes *in vitro*. We then stimulated them with a combination of PRF1 and GZMB at sublytic concentrations^65^ to mimic a cytotoxic T cell attack (Supplementary Fig. 10a). Six hours after stimulation, a dose-dependent increase in F-actin fluorescence was detected (Supplementary Fig. 10be). Live imaging of CellLight Actin-RFP-transfected myelinating ODC confirmed that individual cells showed a relatively stable increase in F-actin levels after stimulation with PRF1 and GZMB (Supplementary Fig. 10c). In addition, pMLC levels increased dose-dependently after stimulation with the cytotoxic molecules (Supplementary Fig. 10d,e). In line with an increased contractile tension of oligodendrocytes after stimulation, MBP^+^ cell surface area was decreased. Strikingly, GZMB + PRF1- dependent F-actin and pMLC induction as well as MBP^+^ surface contraction could be inhibited by simultaneous application of the ROCK inhibitor fasudil (Supplementary Fig. 10f). Our data provide evidence that the immune-driven paranodal constriction of axonal diameters could be mediated by remodeling of the cytoskeleton and actomyosin contractility in myelinating processes.

### Axonal spheroid formation reflects a focal impairment of axonal transport and depends on cytoskeletal dynamics

We and others have previously concluded that axonal spheroids in mice with myelin perturbation form because of focal impairment of axonal transport^26, 66^. To confirm this and identify a putative explanation for pro-degenerative signaling in constricted axons we cross-bred *PLPmut* mice with *Nmnat2*-Venus mice which allows live imaging axonal transport of the survival factor in nerve explants^67^. In 12-month-old mice, the total counts of transported NMNAT2 particles as well as their average and maximum velocities were significantly decreased in optic nerve explants of the myelin mutants (Fig. 8a). In contrast, axons in femoral quadriceps nerve explants of *PLPmut* mice showed similar axonal transport efficacy as in *Wt* mice (Fig. 8b), reflecting the CNS-specific impact of PLP defects in mice. Occasionally, we detected focal accumulations of NMNAT2 particles within individual fibers with little or no ongoing transport in optic nerves of *PLPmut* mice (Fig. 8c). During the 1h live imaging sessions, these accumulations appeared to increase in frequency, indicating ongoing formation of obstructions within damaged fibers in explants. Immunofluorescence identified these accumulations of NMNAT2 to occur within SMI32^+^ axonal spheroids. When analyzing optic nerves from the same mice before and after the imaging, we confirmed that the frequency of SMI32^+^ axonal spheroids was significantly increased after incubating the explants with control medium (DMSO only) (Fig. 8d). In contrast, when incubating the explants with medium containing cytochalasin D or blebbistatin (dissolved in DMSO) to inhibit actin polymerization or myosin activity, respectively, this increase was prevented.

**Fig. 8.**
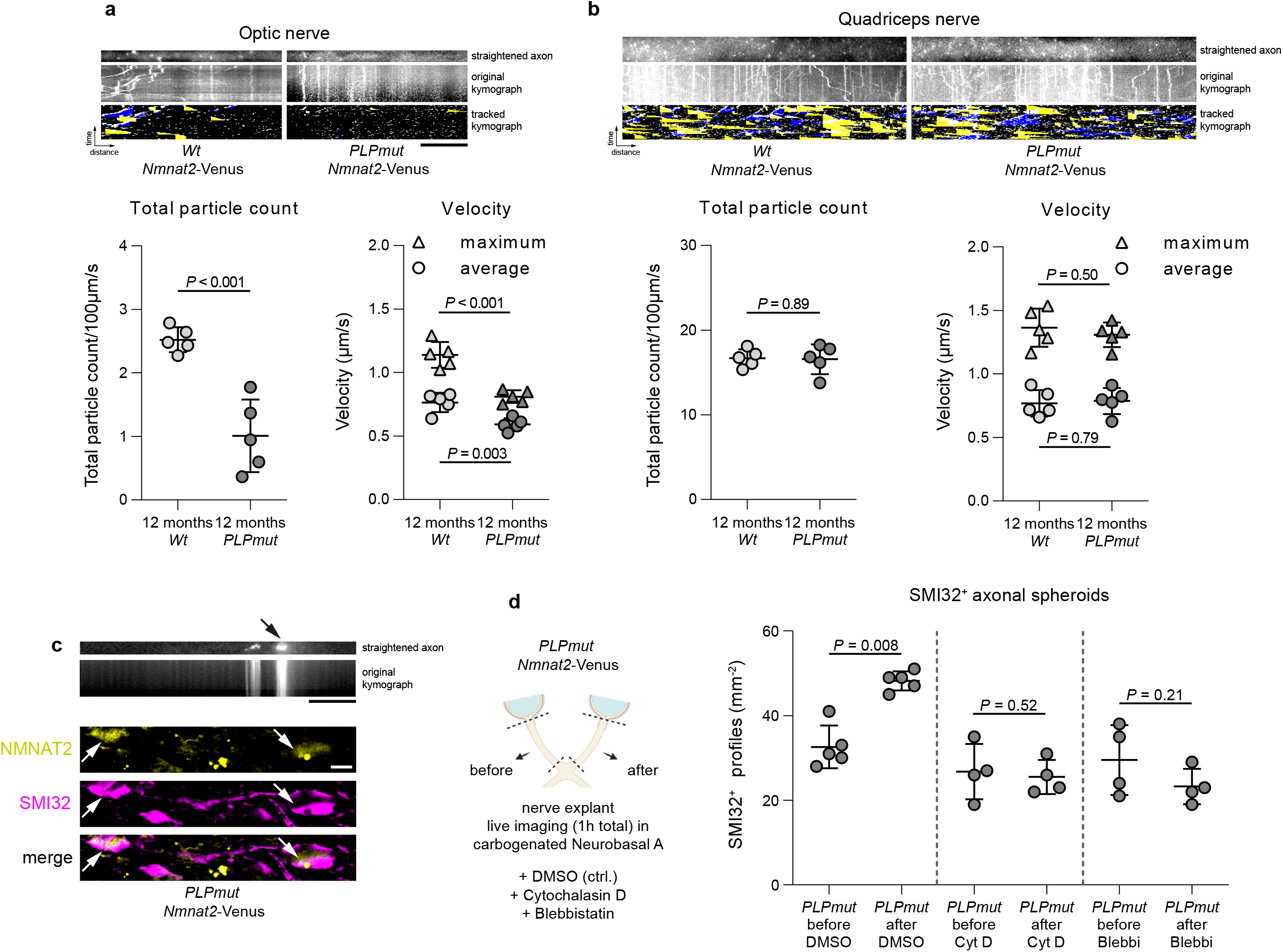
Focal impairment of fast axonal transport in axonal spheroids depends on cytoskeletal plasticity. a. Representative straightened axon, kymograph, and kymograph of successfully tracked particles (top) as well as quantification of axonal transport parameters (bottom) in optic nerve and **b** femoral quadriceps nerve explants from 12-month-old *Wt*/*Nmnat2*-Venus and *PLPmut*/*Nmnat2*-Venus mice. The straightened axon shows the first frame of the time lapse recording (optic nerve: 60 frames in total, quadriceps nerve: 120 frames in total, frame rate 2 fps). Scale bar, 10 µm. Total NMNAT2 particle count and velocity are decreased in CNS but not PNS axons of *PLPmut* mice (each circle represents the mean value of 5 axons of one mouse, *n* = 5 mice per group, two-sided Student’s *t*-test, Optic nerve: Total particle count *t* = 5.602, Maximum velocity *t* = 4.241, Average velocity *t* = 6.478, Quadriceps nerve: Total particle count *t* = 0.1369, Maximum velocity *t* = 0.2749, Average velocity *t* = 0.7069, All d.f. = 8). **c** Straightened axon and kymograph of an optic nerve explant from a *PLPmut*/*Nmnat2*-Venus mouse exemplifying focal accumulation of particles (arrow) in an axon with blocked axonal transport (top). Representative immunofluorescence detection of SMI32 reactivity in optic nerves from 12-month-old *PLPmut*/*Nmnat2*-Venus mice (bottom). The arrows indicate axonal spheroids with an accumulation of NMNAT2 particles. Scale bars, 10 µm. **d** Schematic experimental design (left). SMI32^+^ axonal spheroids were quantified in pairs of optic nerves from *PLPmut*/*Nmnat2*-Venus mice before or after live imaging explants in neurobasal A medium containing DMSO, cytochalasin D (Cyt D), or blebbistatin (Blebbi). Inhibition of actin polymerization or myosin activity blocks the increased frequency of SMI32^+^ axonal spheroids after 1 h imaging *ex vivo* (each circle represents the mean value of one optic nerve, *n* = 4-5 mice per group, two- sided paired *t* test, DMSO: *t* = 4.968, d.f. = 4, Cyt D: *t* = 0.7346, d.f. = 3, Blebbi: *t* = 1.575, d.f. = 3). Data are presented as the mean ± s.d.

We next investigated the consequences of these inhibition experiments on pMLC levels in optic nerve explants. Both cytochalasin D and blebbistatin prevented the increased activation of MLC associated with ongoing axonal spheroid formation in *PLPmut* explants (Fig. 9a). Explants from *Wt* mice did not exhibit a significant increase in spheroid formation during the same imaging timeframe (Fig. 9b), confirming that this ongoing immune-driven constriction process depends on perturbed myelinating glia. As a reciprocal experiment we treated explants with calyculin A, a potent protein phosphatase 1 inhibitor that increases myosin contractility. Moreover, we tested the effect of fasudil on optic nerve explants from *PLPmut* mice. While calyculin A further enhanced pMLC levels and axonal spheroid formation, fasudil inhibited both processes compared with controls (Fig. 9c,d). Finally, we treated *PLPmut* mice for 4 weeks with fasudil in the drinking water to test its efficacy in interfering with ongoing axonal spheroid formation *in vivo*. Importantly, fasudil did not affect the accumulation of CD8^+^ T cells in the white matter of *PLPmut*mice but significantly attenuated the number of axonal spheroids (Fig. 9e), most likely by blocking T cell driven actomyosin constriction at paranodal domains.

**Figure 9.**
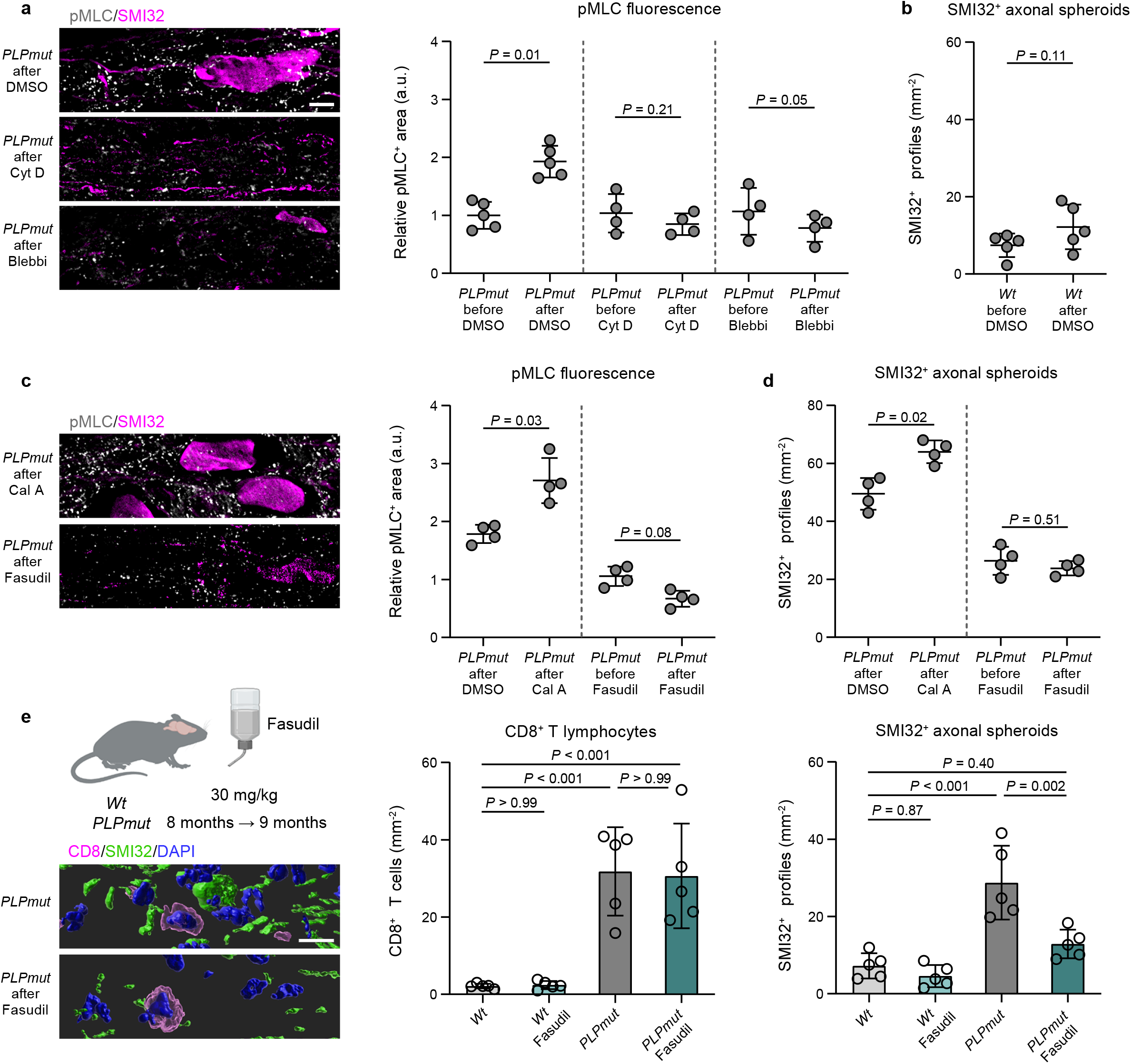
Actomyosin constriction drives axonal spheroid formation in myelin mutant mice. a. Representative immunofluorescence detection (left) of pMLC and SMI32 in optic nerve explants from 12-month-old *PLPmut*/*Nmnat2*-Venus mice after live imaging explants in neurobasal A medium containing DMSO, cytochalasin D (Cyt D), or blebbistatin (Blebbi). Quantification (right) of pMLC fluorescence by thresholding analysis demonstrates an increased relative pMLC^+^ area after live imaging which is blocked by Cyt D or Blebbi (each circle represents the mean value of one optic nerve, *n* = 4-5 mice per group, two-sided paired *t* test, DMSO: *t* = 4.292, d.f. = 4, Cyt D: *t* = 1.613, d.f. = 3, Blebbi: *t* = 3.125, d.f. = 3). **b** Quantification of SMI32^+^ axonal spheroids in pairs of optic nerves from *Wt*/*Nmnat2*-Venus mice before or after live imaging explants in neurobasal A medium containing DMSO. There is no increased frequency of SMI32^+^ axonal spheroids after 1 h imaging *ex vivo* (each circle represents the mean value of one optic nerve, *n* = 5 mice per group, two-sided paired *t* test, *t* = 2.053, d.f. = 4). **c** Representative immunofluorescence detection (left) of pMLC and SMI32 in optic nerve explants from 12-month-old *PLPmut* mice after maintaining explants in neurobasal A medium containing DMSO, calyculin A (Cal A), or fasudil. Quantification (right) of pMLC fluorescence by thresholding analysis and **d** SMI32^+^ axonal spheroids demonstrates a further increase of relative pMLC^+^ area and axonal spheroid formation after Cal A and a block of pMLC induction and axonal spheroid formation after fasudil (each circle represents the mean value of one optic nerve, *n* = 4 mice per group, two-sided paired *t* test, Cal A: *t* = 3.703, d.f. = 3, fasudil: *t* = 2.577, d.f. = 3) **e** Schematic experimental design (left) and IMARIS Z-stack surface rendering of CD8 and SMI32 in the optic nerves of 9-month-old *PLPmut* mice with or without fasudil treatment. Scale bar, 10 µm. Quantification of CD8+ T cells (middle) and SMI32^+^ axonal spheroids (right) demonstrates no impact of fasudil on the numbers of CD8^+^ T cells but on axonal spheroid formation in *PLPmut* mice (each circle represents the mean value of one optic nerve, *n* = 5 mice per group, One-way ANOVA with Tukey’s multiple comparisons test, CD8: *F* (3, 16) = 17.72, *P* < 0.001, SMI32: *F* (3, 16)= 18.99, *P* < 0.001). Data are presented as the mean ± s.d.

In summary, axonal spheroid formation is associated with a focal impairment of fast axonal transport of NMNAT2, which may explain the activation of pro-degenerative signaling typically observed in programmed axon degeneration^68, 69^. Moreover, contractile actomyosin activity (likely occurring in myelinating oligodendrocyte processes) is required for the efficient progression of axonal spheroid formation.

## Discussion

Genetic myelin perturbation can cause glial dysfunction, fiber damage, and functional decline. We have previously discovered that axonal damage is not exclusively a direct consequence of the mutation affecting neuron-glia interactions but that there is an additional impact of neuroinflammation^16, 25, 26^. In line with this, oligodendrocyte-lineage cells are immunocompetent cells able to provide a variety of signals to communicate with innate and adaptive immune cells^12, 14^. Indeed, partially shared disease- and aging-associated reactions of oligodendrocyte-lineage cells have recently been identified and comprise pro-inflammatory molecules^8, 70–72^. Thus, myelin defects seem to result in molecular changes and cell stress pathways in oligodendrocyte subsets that make them susceptible to immune-mediated damage.

Rare genetic disease models display commonalities with more frequent CNS pathologies and offer unique opportunities to investigate the consequences of myelin defects in a defined setting of known etiology and progression. Here we studied such models to identify how axonal damage, myelin loss, and neurodegeneration are related to each other and to disability in the context of detrimental cytotoxic T cell reactions. In both *PLPmut* and *PLPtg* mice, previous studies revealed that CD8^+^ T cells accumulate in the white matter near nodal domains of myelinated axons^26, 28^. While the exact antigen(s) recognized by these cells are not identified, they drive myelin pathology and axonal damage in a cognate TCR-dependent manner using cytotoxic effector molecules^27, 29^. Moreover, a reduction of fast axonal transport efficacy and a focal formation of juxtaparanodal axonal spheroids are shared pathological features in early stages of immune-mediated damage.

In our current study we observed that these spheroids initially form in axonal segments ensheathed by perturbed myelin. Interestingly, axonal spheroids only progress in size and show further signs of axon degeneration when the myelin sheath remains attached. Accordingly, neurodegeneration and clinical disease are inversely related to the degree of demyelination in the distinct disease models. Thus, efficient removal of perturbed myelin segments may allow resilience or recovery of axons at early stages of damage and argues against the current dogma that axon degeneration in myelin disease is mostly secondary to loss of myelin. Nevertheless, there is evidence that the long-term failure to regenerate myelin correlates strongly with neurodegeneration and disability in MS^73–75^. Because previous observations in EAE have shown that focal axonal damage occurs in myelinated fibers and can be reversible^52^, it is important that perturbed myelin is rapidly removed, and replaced by new myelin sheaths. A recent study in human MS and experimental models of inflammatory demyelination is consistent with the notion that myelin itself poses a risk for axon degeneration^76^. Oligodendrocytes under immune attack were proposed to provide insufficient metabolic support to underlying axons and thereby promote their degeneration, which might be reversed by rapid demyelination. Here we describe an additional, mechanical process of how oligodendrocytes targeted by CD8^+^ T cells gain properties detrimental to axonal integrity that could explain focal damage and impairment of axonal transport near the nodal domains.

Myelinating glia in the peripheral nervous system constrict axons after injury to facilitate fragmentation, degradation, and subsequent regeneration^55–57^. This process depends on cytoskeletal dynamics comprising actin-myosin remodeling and is instructed by altered interactions between severed axons and Schwann cells. Recent findings suggest that oligodendrocytes with lowered Dusp6 expression can fulfill similar functions and enable repair after injury^77^. However, other stimuli or defects (e.g., gene defects, cytokines, effector molecules) could induce similar processes in myelinating glia that are to some extent reminiscent of dedifferentiation^78^. Oligodendroglial Vangl2, a RhoA-Myosin II-dependent planar polarity protein, controls paranodal axon diameter^79^, demonstrating some compressive action of the paranodal spiral on the axon. Developmental process extension by oligodendrocytes at the leading edge of myelin sheaths involves actin dynamics^59, 80^. Oligodendrocytes are also known to increase cytoskeletal plasticity after inflammatory damage, which is associated with early changes at the paranodal domains^58, 81^. Our data indicate that increased actomyosin contractility at the paranodal domains can be detrimental to axonal integrity in non-lesioned conditions, which is in line with the vulnerability of fast axonal transport to mechanical constriction^82^. Using various inhibitors to decrease or enhance actomyosin activity in an *ex vivo* explant system and *in vivo*, we demonstrate its involvement in axonal spheroid formation. We do not know where exactly the contractile forces are generated, but there are several findings that point to the paranodal glial domains: First, super-resolution microscopy showed that compact assemblies of F-actin and pMLC form directly adjacent to constricted paranodal axon domains in *PLPmut* mice. Second, a direct mode of action on the neuron is difficult to explain since the membrane periodic skeleton maintains axonal integrity and its disruption via cytoskeletal inhibitors accelerates beading and fragmentation *in vitro*^83, 84^. Third, our experiments using *Wt* explants, as well as the spatiotemporal dynamics of spheroid formation and the above described similarities with myelinating glia in the peripheral nervous system suggest an oligodendrocyte-related mechanism. The predominant localization of cytoskeletal elements within non-compact myelin domains is consistent with the observed paranodal narrowing or “strangulation” of the axonal partners and the accumulation and disintegration of organelles at their juxtaparanodal domains^59^. Nevertheless, we can presently not fully exclude that actomyosin activity within the axon is involved in the constriction. Focal constriction of axonal diameters below a certain threshold could explain the high vulnerability of small diameter myelinated axons in *Plp1* knockout mice which show the same pathology as *PLPmut* mice^26, 85^.

We observed that the combined cytotoxic effector molecules perforin and granzyme B induce the cytoskeletal alterations and the reduction of the myelin membrane sheet surface area in oligodendrocyte cultures, which was blocked by fasudil, an established inhibitor of ROCK-dependent contractility. Moreover, increased pMLC levels in the white matter, paranodal constriction, and axonal spheroid formation in *PLPmut* mice were attenuated by immune deficiency. Since *PLPmut*/*Rag1*^-/-^ mice still exhibit structurally altered myelin, we conclude that these changes are not a cell-intrinsic consequence of the myelin perturbation. Granzyme B can directly cleave kinases and binding proteins important for controlling cytoskeletal dynamics and thereby induce membrane blebbing in target cells^60^. Such plasmalemmal protrusions are also important for physiological processes like cytokinesis and locomotion and can provide resilience against cell lysis and necrosis^86^. It is therefore possible that CD8^+^ T cells directly target myelin segments of perturbed oligodendrocytes and thereby induce cytoskeletal alterations, eventually resulting in abnormal constriction of axons at their paranodal domains. This mechanism could provide an explanation why persistent ensheathment with mutant myelin ultimately results in axonal transection, whereas demyelination abolishes the immune target(s) as well as the constricting oligodendroglial process.

Why oligodendrocytes in the investigated mouse models show distinct reactions upon the CD8^+^ T cell attack remains an open question. While increased expression of markers indicative of disease-associated pro-inflammatory reactions and of ESCRT-mediated membrane repair components is found in both models, upregulation of molecules protecting against phagocytosis and cell death are only found in *PLPmut* mice. This is associated with increased expression of actomyosin-related cytoskeletal molecules and axon degeneration. In contrast, there is increased expression of “eat me” signals in *PLPtg* oligodendrocytes, resulting in the formation of a phagocytic, axon-protective microglia state (AMG2). Similar as in the PNS^87^, the CSF-1-CSF-1R axis seems to play an important part in the communication between neural cells and white matter-resident phagocytes. By modulating microglial myelin phagocytosis, we show that these reactions can be fostered or attenuated, indicating that they are not exclusively driven by oligodendrocyte-intrinsic differences caused by the distinct gene defects. Since both axon degeneration and demyelination are strongly modified by CD8^+^ T cells, these differential glial reactions may instead be induced by distinct subpopulations of CD8^+^ T cells that utilize different effector molecules. In line with this hypothesis, a recent study has shown that aging-related transcriptional responses in oligodendrocytes are at least partially dependent on the presence of CD8^+^ T cells in the white matter^8^. Mild oligodendrocyte cell loss and demyelination was related to an interferon response, while a distinct state of aged oligodendrocytes was characterized by increased expression of *Serpina3n*, an inhibitor of granzyme B. We therefore hypothesize that oligodendrocytes affected by granzyme B try to counteract demyelination and constrict axons, while oligodendrocytes affected by interferon instruct microglia-driven demyelination. This hypothesis is in line with the recently described heterogeneity of CNS-associated CD8^+^ T cells and their effector molecules in myelin mutants and aged mice and is supported by the observation that *Gzmb* deficiency in adaptive immune cells attenuates axonal perturbation but not demyelination in *PLPtg* mice^7, 27, 29^. Moreover, we observed that several transcripts specific for the interferon-responsive oligodendrocyte state (e.g., *Ifi27l2a*, *B2m*) are upregulated in ODC from *PLPtg* but not *PLPmut* mice.

In *PLPtg* mice, microglia driving demyelination (AMG2) showed gene set enrichment for oxidative phosphorylation, typical for tolerogenic antigen presenting cells and phagocytes^88, 89^. Moreover, they upregulated marker genes indicating axon protection and downregulated genes important for T cell stimulation. Thus, they seem to play a beneficial role in removing perturbed myelin and promoting the resolution of inflammation in demyelinated regions which is in line with the preferential accumulation of CD8^+^ T cells in still myelinated regions. However, these microglia also downregulate genes important for lipid export and show accumulation of lysosomal storage material. Thus, overt myelin degradation seems to burden these microglia and might explain the limited remyelination capacity in *PLPtg* mice^90^. On the other hand, a disease-associated microglia state (AMG1) with pro-inflammatory properties was detected in both myelin mutant models, reflecting the context-specific reaction of microglia, and explaining the different outcome after pharmacological microglia depletion (this study and^31^). Partially removing microglia can thus be both beneficial by attenuating neuroinflammatory processes (as observed in *PLPmut* mice) and detrimental by promoting dysfunction of protective reactions (as observed in *PLPtg* mice). Interestingly, it was recently shown that monocyte-derived microglia-like cells with similar properties can infiltrate the brain in aging and disease^43^. However, these DIM show a signature more similar to CAM in our dataset and it was previously demonstrated that there is no significantly increased infiltration of monocyte-derived myeloid cells into the white matter of *PLPtg* mice^91^.

Our observations confirm that focal impairment of axonal transport and formation of axonal spheroids at juxtaparanodal domains are induced by T cell-dependent paranodal alterations. Interestingly, this is associated with activation of pro-degenerative signaling (cleaved caspase3) in axons, reminiscent of trophic deprivation^69^. Since these reactions also occur to some extent in *PLPtg* mice but are less evident in demyelinated regions, they do not necessarily result in axon loss and could be reversible. However, mitochondrial defects and decreased axonal transport of the survival factor NMNAT2 are known to facilitate Wallerian-like axon degeneration^68^. This indicates that later stages of neuroinflammation-driven axonal damage in *PLPmut* mice are likely to be executed by SARM1-dependent, active signaling within axons. While a previous study did not find protection of axons in *Plp1* knockout mice upon crossbreeding with the *Wld*^S^ line^66^, impaired axonal transport could have prevented axonal delivery of the fusion protein. Since we sometimes observed consecutive spheroids along the same axons, we propose that chronic focal damage at multiple myelinated target sites could add up to promote axon degeneration and subsequent neuron loss. Future experiments should aim to clarify the molecular mechanisms of axon degeneration in *PLPmut* mice.

In conclusion, our data show that while myelination offers many advantages for higher functions of the nervous system, it also poses a risk for neuroinflammation-related axonal damage and degeneration when oligodendrocytes become dysfunctional. The evolutionary benefit of myelin appears to outweigh this concomitant risk for insulated axons in pathological conditions. Clearly, demyelination *per se* is generally not beneficial but could provide resilience of compromised fibers allowing survival to possibly enable restoration, as demonstrated in distinct myelin disease models. In this context, a major open question relates to the reason for the susceptibility of myelin perturbation to induce adaptive immune reactions. Oligodendrocytes burdened by myelin defects may incite neuroinflammation as a conserved mechanism to eliminate stressed or dysfunctional cells. Additionally, mutant oligodendrocytes targeted by CD8^+^ T cells might constrict axons by recapitulating a glial program beneficial for clearance of debris and regeneration in injury conditions, similar as Schwann cells during Wallerian degeneration.

Our findings are of translational relevance for therapy approaches to modulate neuroinflammation in myelin disease. While they imply that blocking detrimental immune reactions can have benefit in chronic progressive disease, they also show that certain aspects of neuroinflammation preserve axonal integrity under compromised conditions. Greater insights are needed to identify strategies to block detrimental but still allow or even foster beneficial neural-immune interactions to confer resilience and possibly enable recovery of the perturbed white matter. Moreover, we demonstrate in proof of principle experiments that blocking inflammation-induced contractile forces preserves myelinated axon integrity. Further studies should clarify the efficacy of such therapeutic approaches in more detail.

## Methods

### Animals

Mice were kept at the animal facility of the Centre for Experimental Molecular Medicine, University of Würzburg, under barrier conditions and at a constant cycle of 14 h in the light (<300 lux) and 10 h in the dark. Colonies were maintained at 20-24 °C and 40-60% humidity, with free access to food and water. All animal experiments were approved by the Government of Lower Franconia, Germany. All mice including wild type (*Wt*), homozygous *PLPmut* (R137W, B6.Cg-Tg(PLP1)1Rm*-Plp1^tm1Kan^*/J)^26^ - genuine or crossbred with *Rag1*^-/-^ (B6.129S7-*Rag1^tm1Mom^*/J)^92^, hemizygous *PLPtg* (B6.Cg-Tg(Plp1)66Kan/J)^34^, mice were on a uniform C57BL/6J genetic background; they were bred, regularly backcrossed and aged in-house. *Nmnat2-*Venus^93^ mice were cross-bred with *PLPmut* mice. *Wt* and *PLPmut* offspring hemizygous for the *Nmnat2* transgene were analyzed. Since we did not detect obvious differences between male and female mice in the analyses presented in the current study, mice of either sex were used for the experiments. Genotypes were determined by conventional PCR using isolated DNA from ear punch biopsies according to previously published protocols.

### Accelerating rotarod analysis

Mice were placed on a RotaRod Advanced system (TSE systems); the time on the constantly accelerating rod (5-50 r.p.m., max latency 300 s) was measured in 5 consecutive runs per trial as described previously^26^. Mice were trained with two trials on two consecutive days and latencies were measured in a third trial on the third day.

### Analysis of visual acuity

The visual acuity of mice was analyzed using automated optokinetic reflex tracking in an OptoDrum device (Striatech). Briefly, mice were placed on an elevated platform surrounded by monitors and a stripe pattern with maximum contrast and constant rotation speed (12 deg s^−1^) was presented. Behavior was automatically detected and analyzed by the OptoDrum software v.1.2.6 in an unbiased manner and the stimulus pattern (cycles) was continuously adjusted to find the threshold of the animal’s visual acuity.

### Spectral domain optical coherence tomography (OCT)

Mice were subjected to OCT imaging with a commercially available device (SPECTRALIS OCT; Heidelberg Engineering) and additional lenses as described previously^26^. Mice were measured at different ages for longitudinal analysis and the thickness of the innermost retinal composite layer comprising the nerve fiber layer (NFL), GCL and inner plexiform layer (IPL) were measured in high-resolution peripapillary circle scans (at least ten measurements per scan) by an investigator unaware of the genotype and treatment condition of the mice using HEYEX v.1.7.1.

### Histochemistry and immunofluorescence

Mice were euthanized with CO_2_ (according to the guidelines by the State Office of Health and Social Affairs Berlin), blood was removed by transcardial perfusion with PBS containing heparin and tissue was fixed by perfusion with 2% paraformaldehyde (PFA) in PBS. Tissue was collected, postfixed, dehydrated, and processed as described previously^26^. Immunohistochemistry was performed on 10- or 30-μm thick longitudinal optic nerve and coronal brain sections after postfixation in 4% PFA in PBS or ice-cold acetone for 10 min. Sections were blocked using 5% BSA in PBS and incubated overnight at 4 °C with an appropriate combination of up to 3 of the following antibodies or stains: mouse anti-neurofilament H non-phosphorylated, SMI32 (1:1,000, catalog no. 801701; BioLegend); rat anti-CD8 (1:500, catalog no. MCA609G; Bio-Rad Laboratories); rabbit anti-MBP (1:300, catalog no. PD004, MBL); rat anti-CD11b (1:100, catalog no. MCA74G; Bio-Rad Laboratories); hamster anti- CD11c (1:100, catalog no. MA11C5; Thermo Fisher Scientific); rabbit anti-P2RY12 (1:300, catalog no. 55043A, AnaSpec); rabbit anti-GAL3 (1:1,000, NBP3-03252; Novus Biologicals); rat anti-GAL3 (1:300, catalog no. 125402, BioLegend); phalloidin-TRITC (1:300, catalog no. P1951; Sigma-Aldrich); FluoroMyelin Green (1:300, catalog no. F34651, Thermo Fisher Scientific); rabbit anti-cleaved caspase 3 (1:300, catalog no. 9664; Cell Signaling); rabbit anti-pMLC Thr18/Ser19 (1:100, catalog no. 3674; Cell Signaling); mouse anti-CASPR (1:1000, catalog no. 75-001; NeuroMab); goat anti-IBA1 (1:300, catalog no. NB100-1028; Novus Biologicals); goat anti-ICAM1(1:1,000, catalog no. AF796; Novus Biologicals); rabbit anti-laminin (1:300, catalog no. ab11575; Abcam), rabbit anti-c-Jun (1:300, catalog no. 9165; Cell Signaling); rat anti-MHC-I (1:100, catalog no. T-2105; Dianova); Immunoreactive profiles were visualized using fluorescently labeled (1:300; Dianova) secondary antibodies, streptavidin (1:300; Thermo Fisher Scientific) or biotinylated secondary antibodies (1:100; Vector Laboratories) and streptavidin-biotin-peroxidase (Vector Laboratories) complex using diaminobenzidine HCl and H_2_O_2_; nuclei were stained with 4,6-diamidino-2-phenylindole (DAPI) (Sigma-Aldrich). Light and fluorescence microscopy images were acquired using an Axio Imager M2 microscope (ZEISS) with ApoTome.2 structured illumination equipment, attached Axiocam cameras and corresponding software (ZEN v.2.3 blue edition) or a FluoView FV1000 confocal microscope (Olympus) with corresponding software (v.2.0). Images were minimally processed (rotation, cropping, addition of symbols) to generate figures using Photoshop CS6 and Illustrator CS6 (Adobe). Z-stack surface rendering was performed using IMARIS v.9.7 (Bitplane). For quantification, immunoreactive profiles were counted in at least three nonadjacent sections for each animal and related to the area of these sections using the cell counter plugin in Fiji/ImageJ v.1.51 (National Institutes of Health). To quantify RGC, perfusion-fixed eyes were enucleated, and specific markers of the inner retinal cell types were labeled in free-floating retina preparations. Fixed retinae were frozen in PBS containing 2% Triton X-100, thawed, washed, and blocked for 1 h using 5% BSA and 5% donkey serum in PBS containing 2% Triton X-100. Retinae were incubated overnight on a rocker at 4 °C with appropriate combinations of the following antibodies: guinea pig anti-RBPMS (1:300, catalog no. ABN1376; Merck Millipore); goat anti-Brn3a (1:100, catalog no. sc- 31984; Santa Cruz Biotechnology); immune reactions were visualized using fluorescently labeled (1:500; Dianova) secondary antibodies, retinae were flat- mounted, and the total retinal area was measured. RGC were quantified in three images of the middle retinal region per flat mount using the cell counter plugin in Fiji/ImageJ v.1.51 (National Institutes of Health).

### Electron microscopy

The optic nerves of transcardially perfused mice were postfixed overnight in 4% PFA and 2% glutaraldehyde in cacodylate buffer. Nerves were osmicated and processed for light and electron microscopy; morphometric quantification of neuropathological alterations was performed as published previously^26^ using a LEO906 E electron microscope (ZEISS) and corresponding software iTEM v.5.1 (Soft Imaging System). At least 10 regions of interest (corresponding to an area of around 5% and up to 3,000 axons per individual optic nerve) were analyzed per optic nerve per mouse. The percentages of axonal profiles showing spheroid formation or undergoing degeneration were identified individually by their characteristic morphological features in electron micrographs and related to the number of all investigated axons per optic nerve per mouse. Images were processed (rotation, cropping, addition of symbols and pseudocolor) to generate figures using Photoshop CS6 and Illustrator CS6 (Adobe).

### Flow cytometry and cell sorting

Mice were euthanized with CO_2_ (according to the guidelines by the State Office of Health and Social Affairs Berlin) and blood was thoroughly removed by transcardial perfusion with PBS containing heparin. Brains including optic nerves, leptomeninges and choroid plexus were dissected, collected in ice-cold PBS and cut into small pieces. Tissue was digested in 1 ml of Accutase (Merck Millipore) per brain at room temperature for 15 min and triturated through 70-μm cell strainers, which were rinsed with 10% FCS in PBS. Cells were purified by a linear 40% Percoll (GE Healthcare) centrifugation step at 650 g without brakes for 25 min and the myelin top layer and supernatant were discarded. Mononuclear cells were resuspended in 1% BSA in PBS and isolated cells were counted for each brain. For scRNA-seq, cells from the brains of 3 adult (10-month-old) *Wt*, 3 *PLPmut*, and 3 *PLPtg* mice were analyzed. Viable cells were identified by Calcein blue AM stain (catalog no. ABD-22007; Biomol), Fc receptors were blocked for 15 min with rat anti-CD16/32 (1:200, catalog no. 553141; BD Biosciences) and cells were washed and labeled with the following antibodies for 30 min at 4 °C: rat anti-CD45 PerCP/Cyanine5.5 (1:100, catalog no. 130-102-469; Miltenyi Biotec); rat anti-SiglecH PE (1:100, catalog no. 12-0333-82; eBioscience); rat anti-O1 AF700 (1:100, FAB1327N-100UG; R&D Systems). Cells were washed twice, single viable cells were gated and CD45^-^O1^+^ and CD45^low^Siglec- H^+^ cells were collected using a FACSAria III and corresponding software (FACSDiva, v.6; BD Biosciences). Calculation of the number of CD45^low^SiglecH^+^ microglia per brain was performed by extrapolating their frequency to the counted total number of isolated cells. For further experiments viable CD45^low^Siglec-H^+^ microglia were labeled with rat anti-ICAM1 BV711 (1:100, catalog no. 116143; BioLegend), rat anti- Ly6A/E FITC (1:100, catalog no. 108106; BioLegend), rat anti-CD11c APC (1:100, catalog no. 117310: BioLegend), and rat anti-PD1 BV605 (1:100, catalog no. 135219; BioLegend). Cells were washed twice; single viable cells were gated and CD45^low^Siglec-H^+^ cells were analyzed using a FACSLyric (BD Biosciences) and Flowjo (version 10).

### Single-cell RNA sequencing (scRNA-seq) and data processing

Around 15,000 CD45^low^Siglec-H^+^ and CD45^-^O1^+^ single cells each were sorted per sample using a FACSAria III (BD Biosciences) before being encapsulated into droplets with the Chromium Controller (10x Genomics) and processed according to the manufacturer’s specifications. Briefly, every transcript captured in all the cells encapsulated with a bead was uniquely barcoded using a combination of a 16-base pair (bp) 10x barcode and a 10-bp unique molecular identifier (UMI). Complementary DNA libraries ready for sequencing on Illumina platforms were generated using the Chromium Single Cell 3′ Library & Gel Bead Kit v2 (10x Genomics) according to the detailed protocol provided by the manufacturer. Libraries were quantified by Qubit 3.0 Fluorometer (Thermo Fisher Scientific) and quality was checked using a 2100 Bioanalyzer with High Sensitivity DNA kit (Agilent Technologies). Libraries were pooled and sequenced with a NovaSeq 6000 platform (S1 Cartridge; Illumina) in paired-end mode to reach a mean of 44,131 reads per single cell. A total of 13,556, 9,955, and 12,191 cells were captured and a median gene number per cell of 2,268, 2,562, and 2,165 could be retrieved for adult *Wt*, *PLPmut*, and *PLPtg* cells, respectively. Data were demultiplexed using the CellRanger software v.2.0.2 based on 8 bp 10x sample indexes; paired-end FASTQ files were generated. The cell barcodes and transcript unique molecular identifiers were processed as described previously^94^. The reads were aligned to the University of California, Santa Cruz mouse mm10 reference genome using STAR aligner v.2.5.1b. The alignment results were used to quantify the expression level of mouse genes and generate the gene-barcode matrix. Subsequent data analysis was performed using the R package Seurat^95^ v.4.0. Doublets and potentially dead cells were removed based on the percentage of mitochondrial genes (cutoff set at 60%) and the number of genes (cells with >100 and <6,000 genes were used) expressed in each cell as quality control markers. The gene expression of the remaining cells (13,522, 9,898, and 12,170 cells from *Wt*, *PLPmut*, and *PLPtg* mice, respectively) was log-normalized. Highly variable genes were detected with Seurat and the top 2,000 of these genes were used as the basis for downstream clustering analysis. Data were scaled, principle component analysis was used for dimensionality reduction and the number of principal components was identified using the built-in Elbow plot function. Cells were clustered based on the identified principal components (10) with a resolution of 0.5; uniform manifold approximation and projection was used for data visualization in two dimensions. Microglia and oligodendrocyte clusters were subset based on marker gene expression and reanalyzed, resulting in 9,484, 8,405, and 8,419 cells from *Wt*, *PLPmut*, and *PLPtg* mice, respectively. Due to their vulnerability during the processing after sorting, much lower numbers of oligodendrocytes compared with microglia remained in the resulting dataset. These were reanalyzed and reclustered based on 14 principal components and a resolution of 0.6. Contribution of the samples to each microglia cluster in absolute numbers was calculated by extrapolating their frequencies to the number of CD45^low^Siglec-H^+^ cells per brain.

Differentially expressed genes were identified using the FindMarkers function with min.pct = 0.25. Complete lists of differentially expressed genes (p_val_adj > 0.05 after Bonferroni or FDR correction) are included in Supplementary Table 1. Marker gene scores for feature expression programs were calculated using the AddModuleScore function in Seurat. Gene-set enrichment analysis was performed using Metascape v.3.5 (http://metascape.org)^96^.

### Cuprizone treatment

Demyelination was induced by feeding 10-week-old *Wt* and *PLPmut* mice a diet containing 0.2% cuprizone (bis-cyclohexanone oxaldihydrazone; Sigma-Aldrich) in ground standard rodent chow for 6 weeks. Control mice were fed standard rodent chow. Optic nerves and corpora callosa were processed for immunofluorescence and electron microscopy. The procedures were approved by the Review Board for the Care of Animal Subjects of the district government (Niedersächsisches Landesamt für Verbraucherschutz und Lebensmittelsicherheit, Germany, AZ 15/1762) and performed according to international guidelines on the use of laboratory mice.

### PLX5622 treatment

PLX5622 (provided by Plexxikon Inc., Berkeley, CA, USA) was prepared as a 300 ppm drug chow to dose ∼L54 mg PLX5622/kg body weight when given *ad libitum*. This was based on our previous long-term treatment approaches in which we observed efficient microglia depletion without obvious neurological side effects in *Wt* and *PLPmut* mice^31^. Control mice received normal chow without the pharmacological inhibitor. *Wt* and *PLPtg* mice were treated for 6 months with daily monitoring concerning certain burden criteria and phenotypic abnormalities. The treatment started at 6 months of age before prominent demyelination in *PLPtg* mice.

### RNAscope

Multiplex RNAscope was performed based on the manufacturer’s (Advanced Cell Diagnostics) instructions^97^. Briefly, sections were postfixed in ice-cold 4% PFA for 30 min. Sections were then dehydrated using a series of ethanol solutions (50 - 100%) before drying and boiling for 5 min in fresh target retrieval solution at 100 °C. Slides were washed in distilled water and rinsed with 100% ethanol before incubation with protease III for 30 min at 40 °C. Slides were washed again and hybridized with gene-specific probes against *Csf1* and *Mbp* for 2 h at 40 °C in a HybEZ oven (ACD). Non-annealed probes were removed by washing sections in 1x proprietary wash buffer, and the signal was amplified with 4 amplification systems (Amp1 - Amp4). Finally, sections were stained with DAPI, mounted and analyzed using an Axio Imager M2 microscope (ZEISS) with ApoTome.2 structured illumination equipment, attached Axiocam cameras and corresponding software (ZEN v.2.3 blue edition).

### Expansion microscopy

Super-resolution fluorescence microscopy was performed with 30-μm-thick longitudinal optic nerve cryo-sections. Free-floating sections were blocked using 5% BSA and 5% donkey serum in PBS containing 0.3% Triton X-100 and incubated overnight at 4 °C with a combination of the following antibodies and stains: rabbit anti-pMLC Thr18/Ser19 (1:100, catalog no. 3674; Cell Signaling); mouse anti-CASPR (1:1000, catalog no. 75-001; NeuroMab); biotin-XX phalloidin (1:300, catalog no. sc-505886; Santa Cruz Biotechnology); Labeling was visualized using AF555 donkey anti-rabbit (1:300; Dianova), CF640R donkey anti-mouse (1:300; Dianova), and streptavidin AF488 (1:300; Thermo Fisher Scientific). Proteins were anchored using 0.1 mg/ml acryloyl-X (AcX, catalog no. A20770; Thermo Fisher Scientific) in PBS for 24 h at room temperature. Sections were washed, partially air-dried on an uncharged slide, incubated in gelling solution [8.6 g/100 ml sodium acrylate (Sigma-Aldrich, catalog no. 408220), 2.5 g/100 ml acrylamide (Sigma-Aldrich, catalog no. A8887), 0.1 g/100 ml N,N′-methylenebisacrylamide (Sigma-Aldrich, catalog no. M7279), 11.7 g/100 ml Sodium chloride (Sigma-Aldrich, catalog no. S6191), 0.2% TEMED accelerator solution (Sigma-Aldrich, catalog no. T9281), 0.01% 4-hydroxy-TEMPO inhibitor solution (Sigma-Aldrich, catalog no. 176141), 0.2% ammonium persulfate (Sigma Aldrich, catalog no. 248614)] for 1 h at 4 °C, and embedded in fresh gelling solution in an assembled chamber using coverslips as spacers and cover. Polymerization was performed at 37 °C for 2 h. Tissue-containing gels were trimmed, scooped off the slide, and digested in Proteinase K overnight at room temperature. Nuclei were labelled with DAPI, and gels were expanded by repeated washes in distilled water until expansion plateaued. Post-expansion specimens were imaged using a LSM900 confocal microscope (Zeiss) with an AiryScan2 detector using an LD-C Apochromat 40x/1.1 W objective. The expansion factor was calculated by measuring gels and labeled structures before and after expansion.

### Primary oligodendrocyte cell cultures

OPCs were prepared from P7 C57BL/6J mouse brains by immunopanning^98^. Briefly, brains were digested with papain and dissociated to single-cell suspension, which was passed through two negative-selection plates coated with BSL1 to remove microglia. The remaining cell suspension was then incubated in a positive-selection plate coated with anti-PDGFRα antibodies (catalog no. sc-338; Santa Cruz Biotechnology). The attached cells were collected using trypsin and cultured on poly(L-lysine)-coated coverslips in proliferation medium containing Dulbecco’s modified Eagle’s medium (DMEM, catalog no. 41965; Thermo Fisher Scientific), Sato Supplement, B-27 Supplement, GlutaMAX, Trace Elements B, penicillin– streptomycin, sodium pyruvate, insulin, N-acetyl-L-cysteine, D-biotin, forskolin, ciliary neurotrophic factor (CNTF), platelet-derived growth factor (PDGF) and neurotrophin- 3 (NT-3). OPCs were cultured in differentiation medium containing DMEM (Thermo Fisher Scientific), Sato Supplement, B-27 Supplement, GlutaMAX, Trace Elements B, penicillin–streptomycin, sodium pyruvate, insulin, N-acetyl-l-cysteine, D-biotin, forskolin, CNTF and NT-3. After 7 days, when OPCs had differentiated into oligodendrocytes, they were stimulated with medium containing sublytic^65^ final concentrations (0.2 or 0.4 µg/ml) of PRF1 (catalog no. APB317Mu01; Cloud-Clone Corp.) + GZMB (catalog no. ab50114; Abcam). For experiments to inhibit ROCK-dependent actomyosin contractility, fasudil (20 µM, catalog no. orb746457; Biorbyt) was applied simultaneously. After 6 hours, the cultures were fixed and analyzed by immunocytochemistry. For live imaging, oligodendrocytes were transfected with CellLight Actin-RFP (catalog no. C10502; Thermo Fisher Scientific) and imaged before and after stimulation using a Leica DMi8 microscope equipped with the DMC 2900/DFC 3000 G camera control, a stage top incubation system (Ibidi), and LAS X software (Leica).

### Live imaging of axonal transport in nerve explants

Fast axonal transport was analyzed in optic nerves and femoral quadriceps nerves of *Wt* and *PLPmut* mice hemizygous for the NMNAT2-Venus transgene as previously described^67^. Briefly, nerves were rapidly dissected into prewarmed (37°C), pre-oxygenated Neurobasal-A medium (Gibco). Imaging of axonal transport in tissue explants was performed using a Leica DMi8 microscope equipped with the DMC 2900/DFC 3000 G camera control, a stage top incubation system (ibidi), and LAS X software (Leica). During imaging, nerves were maintained in oxygenated Neurobasal-A medium at 37°C. Images were captured using fixed light intensity and camera exposure time settings at a rate of 2 frames per second and 60 frames over a total imaging period of 1 hour. Afterwards, the nerves were fixed and analyzed by immunocytochemistry. At least 3 individual movies (often containing multiple axons) were captured for each nerve explant. Individual axons were straightened using the Straighten plugin in Fiji/ImageJ v.1.51 (National Institutes of Health). Axonal transport parameters were determined for individual axons using the DifferenceTracker^99^ set of plugins using previously described parameters^67^. On average, 5 axons were analyzed for each nerve explant. For inhibition/stimulation of actomyosin contractility, nerves were incubated with medium containing DMSO (control), cytochalasin D (1 µg/ml, catalog no. C6637; Sigma-Aldrich), blebbistatin (100 µM, catalog no. B0560; Sigma-Aldrich), calyculin A (0.1 µM, catalog no. C5552; Sigma-Aldrich), or fasudil (20 µM, catalog no. orb746457; Biorbyt) and compared with nerves from the same animal fixed before and after the 1 h imaging session.

### Fasudil treatment

Fasudil (catalog no. orb746457; Biorbyt) was prepared at 180 µg/ml within the drinking water to dose ∼L30 mg/kg body weight per day when given *ad libitum*. This was based on previous long-term treatment approaches *in vivo* which achieved efficient attenuation of brain pathology in distinct disease models^100, 101^. Control mice received normal drinking water without the pharmacological inhibitor. *Wt* and *PLPmut* mice were treated for 4 weeks with daily monitoring concerning certain burden criteria and phenotypic abnormalities. The treatment started at 8 months of age when neuroinflammation and axon degeneration are ongoing in *PLPmut* mice.

### Statistics and reproducibility

All quantifications and analyses were performed by blinded investigators who were unaware of the genotype and treatment group of the respective mice or tissue samples after concealment of genotypes with individual uniquely coded labels. Animals were randomly placed into experimental or control groups according to the genotyping results using a random generator (http://www.randomizer.org). For biometrical sample size estimation, G*Power v.3.1.3 was used^102^. Calculation of appropriate sample size groups was performed using an *a priori* power analysis by comparing the mean of 2 to 3 groups with a defined adequate power of 0.8 (1 - beta error) and an α error of 0.05. To determine the prespecified effect size d or f, previously published data were considered as comparable reference values^26^. The number of individual mice per group (number of biologically independent samples) for each experiment and the meaning of each data point are indicated in the respective figure legends. All data (except scRNA-seq) represent at least three independent experiments. For the histological analyses, we quantified a specific cell type/structure in at least three different sections of a respective tissue and averaged the measurements into one single data point. No animals were excluded from the analyses. In the scRNA-seq experiment, we analyzed the brains of 3 mice for each group. Statistical analysis was performed using Prism 8 (GraphPad Software). The Shapiro-Wilk test was used to check for the normal distribution of data and the *F* test was used to check the equality of variances to ensure that all data met the assumptions of the statistical tests used. Comparisons of two groups were performed with an unpaired Student’s t-test (parametric comparison) or Mann-Whitney *U*-test (nonparametric comparison). For multiple comparisons, a one-way analysis of variance (ANOVA) (parametric) or Kruskal-Wallis test (nonparametric) with Tukey’s post hoc test were applied and adjusted *P* values are presented. *P* < 0.05 was considered statistically significant; exact *P* values are provided whenever possible in the figures and/or figure legends.

## Data availability

The sequencing data are available from the Gene Expression Omnibus. Other data that support the findings of this study are available from the corresponding authors.

## Acknowledgements

We thank H. Blazyca and B. Meyer for technical assistance and J. Schreiber, A. Weidner, and T. Bimmerlein for their attentive care of mice. We are grateful to D. Klein for valuable discussions. This work was supported by the German Research Foundation (grant no. MA1053/6-2 to R.M. and grant no. GR5240/1-1 to J.G.), the Roman, Marga und Mareille Sobek Foundation (to R.M.), the Charitable Hertie Foundation (grant no. P1150084 to J.G.) and the Interdisciplinary Centre for Clinical Research of the University of Würzburg (grant no. A-302 to R.M.). We thank Plexxikon for generous support and providing PLX5622.

## Author contributions

J.G. and R.M. planned and oversaw all aspects of the study. J.G., T.A., Y.K., and M.H. performed and analyzed most of the experiments. S.L. performed isolation and culture of oligodendrocyte-lineage cells. V.G. and M.S. (Hannover) performed the cuprizone treatment experiment. R.A. and M.C. provided *Nmnat2*-Venus mice and instructions for analysis of fast axonal transport. F.I. and A.E.S. performed cell sorting and scRNA-seq. M. S. (Munich) provided equipment and advice for super- resolution microscopy and inhibition experiments. J.G. wrote the manuscript with input from all authors.

## References

1. Simons, M. & Nave, K. A. Oligodendrocytes: Myelination and Axonal Support. Cold Spring Harb Perspect Biol 8, a020479 (2015).

2. Bonetto, G., Belin, D. & Karadottir, R. T. Myelin: A gatekeeper of activity-dependent circuit plasticity? Science 374, eaba6905 (2021).

3. Duncan, I. D. & Radcliff, A. B. Inherited and acquired disorders of myelin: The underlying myelin pathology. Exp Neurol 283, 452–475 (2016).

4. Salvadores, N., Sanhueza, M., Manque, P. & Court, F. A. Axonal Degeneration during Aging and Its Functional Role in Neurodegenerative Disorders. Front Neurosci 11, 451 (2017).

5. Stassart, R. M., Möbius, W., Nave, K. A. & Edgar, J. M. The Axon-Myelin Unit in Development and Degenerative Disease. Front Neurosci 12, 467 (2018).

6. Filley, C. M. White matter and human behavior. Science 372, 1265–1266 (2021).

7. Groh, J., et al. Accumulation of cytotoxic T cells in the aged CNS leads to axon degeneration and contributes to cognitive and motor decline. Nature Aging 1, 357–367 (2021).

8. Kaya, T., et al. CD8(+) T cells induce interferon-responsive oligodendrocytes and microglia in white matter aging. Nat Neurosci (2022).

9. Schetters, S. T. T., Gomez-Nicola, D., Garcia-Vallejo, J. J. & Van Kooyk, Y. Neuroinflammation: Microglia and T Cells Get Ready to Tango. Front Immunol 8, 1905 (2017).

10. Matejuk, A., Vandenbark, A. A. & Offner, H. Cross-Talk of the CNS With Immune Cells and Functions in Health and Disease. Front Neurol 12, 672455 (2021).

11. Yong, V. W. Microglia in multiple sclerosis: Protectors turn destroyers. Neuron (2022).

12. Peferoen, L., Kipp, M., van der Valk, P., van Noort, J. M. & Amor, S. Oligodendrocyte-microglia cross-talk in the central nervous system. Immunology 141, 302–313 (2014).

13. Falcao, A. M., et al. Disease-specific oligodendrocyte lineage cells arise in multiple sclerosis. Nat Med 24, 1837–1844 (2018).

14. Kirby, L. & Castelo-Branco, G. Crossing boundaries: Interplay between the immune system and oligodendrocyte lineage cells. Semin Cell Dev Biol 116, 45–52 (2021).

15. Absinta, M., et al. A lymphocyte-microglia-astrocyte axis in chronic active multiple sclerosis. Nature 597, 709–714 (2021).

16. Groh, J. & Martini, R. Neuroinflammation as modifier of genetically caused neurological disorders of the central nervous system: Understanding pathogenesis and chances for treatment. Glia 65, 1407–1422 (2017).

17. Chen, J. F., et al. Enhancing myelin renewal reverses cognitive dysfunction in a murine model of Alzheimer’s disease. Neuron 109, 2292-2307 e2295 (2021).

18. Tansey, M. G., et al. Inflammation and immune dysfunction in Parkinson disease. Nat Rev Immunol (2022).

19. Trapp, B. D. & Stys, P. K. Virtual hypoxia and chronic necrosis of demyelinated axons in multiple sclerosis. Lancet Neurol 8, 280–291 (2009).

20. Friese, M. A., Schattling, B. & Fugger, L. Mechanisms of neurodegeneration and axonal dysfunction in multiple sclerosis. Nat Rev Neurol 10, 225–238 (2014).

21. Duncan, G. J., Simkins, T. J. & Emery, B. Neuron-Oligodendrocyte Interactions in the Structure and Integrity of Axons. Front Cell Dev Biol 9, 653101 (2021).

22. Trapp, B. D. & Nave, K. A. Multiple sclerosis: an immune or neurodegenerative disorder? Annu Rev Neurosci 31, 247–269 (2008).

23. Edgar, J. M., et al. Demyelination and axonal preservation in a transgenic mouse model of Pelizaeus-Merzbacher disease. EMBO Mol Med 2, 42–50 (2010).

24. Smith, C. M., Cooksey, E. & Duncan, I. D. Myelin loss does not lead to axonal degeneration in a long-lived model of chronic demyelination. J Neurosci 33, 2718-2727 (2013).

25. Ip, C. W., et al. Immune cells contribute to myelin degeneration and axonopathic changes in mice overexpressing proteolipid protein in oligodendrocytes. J Neurosci 26, 8206–8216 (2006).

26. Groh, J., et al. Pathogenic inflammation in the CNS of mice carrying human PLP1 mutations. Hum Mol Genet 25, 4686–4702 (2016).

27. Kroner, A., Ip, C. W., Thalhammer, J., Nave, K. A. & Martini, R. Ectopic T-cell specificity and absence of perforin and granzyme B alleviate neural damage in oligodendrocyte mutant mice. Am J Pathol 176, 549–555 (2010).

28. Ip, C. W., et al. Neuroinflammation by cytotoxic T-lymphocytes impairs retrograde axonal transport in an oligodendrocyte mutant mouse. PLoS One 7, e42554 (2012).

29. Abdelwahab, T., et al. Cytotoxic CNS-associated T cells drive axon degeneration by targeting perturbed oligodendrocytes in PLP1 mutant mice. iScience 26, 106698 (2023).

30. Ip, C. W., Kroner, A., Crocker, P. R., Nave, K. A. & Martini, R. Sialoadhesin deficiency ameliorates myelin degeneration and axonopathic changes in the CNS of PLP overexpressing mice. Neurobiol Dis 25, 105–111 (2007).

31. Groh, J., Klein, D., Berve, K., West, B. L. & Martini, R. Targeting microglia attenuates neuroinflammation-related neural damage in mice carrying human PLP1 mutations. Glia 67, 277–290 (2019).

32. Anderson, T. J., et al. Late-onset neurodegeneration in mice with increased dosage of the proteolipid protein gene. J Comp Neurol 394, 506–519 (1998).

33. Groh, J., Horner, M. & Martini, R. Teriflunomide attenuates neuroinflammation-related neural damage in mice carrying human PLP1 mutations. J Neuroinflammation 15, 194 (2018).

34. Readhead, C., Schneider, A., Griffiths, I. & Nave, K. A. Premature arrest of myelin formation in transgenic mice with increased proteolipid protein gene dosage. Neuron 12, 583–595 (1994).

35. Masuda, T., Sankowski, R., Staszewski, O. & Prinz, M. Microglia Heterogeneity in the Single-Cell Era. Cell Rep 30, 1271–1281 (2020).

36. Bisht, K., et al. Capillary-associated microglia regulate vascular structure and function through PANX1-P2RY12 coupling in mice. Nat Commun 12, 5289 (2021).

37. Sala Frigerio, C., et al. The Major Risk Factors for Alzheimer’s Disease: Age, Sex, and Genes Modulate the Microglia Response to Abeta Plaques. Cell Rep 27, 1293–1306 e1296 (2019).

38. Keren-Shaul, H., et al. A Unique Microglia Type Associated with Restricting Development of Alzheimer’s Disease. Cell 169, 1276–1290 e1217 (2017).

39. Krasemann, S., et al. The TREM2-APOE Pathway Drives the Transcriptional Phenotype of Dysfunctional Microglia in Neurodegenerative Diseases. Immunity 47, 566–581 e569 (2017).

40. Safaiyan, S., et al. White matter aging drives microglial diversity. Neuron 109, 1100–1117 e1110 (2021).

41. Jordao, M. J. C., et al. Single-cell profiling identifies myeloid cell subsets with distinct fates during neuroinflammation. Science 363 (2019).

42. Sousa, C., et al. Single-cell transcriptomics reveals distinct inflammation-induced microglia signatures. EMBO Rep 19 (2018).

43. Silvin, A., et al. Dual ontogeny of disease-associated microglia and disease inflammatory macrophages in aging and neurodegeneration. Immunity 55, 1448–1465 e1446 (2022).

44. Deczkowska, A., et al. Disease-Associated Microglia: A Universal Immune Sensor of Neurodegeneration. Cell 173, 1073–1081 (2018).

45. Goldmann, T., et al. USP18 lack in microglia causes destructive interferonopathy of the mouse brain. EMBO J 34, 1612–1629 (2015).

46. Hasel, P., Rose, I. V. L., Sadick, J. S., Kim, R. D. & Liddelow, S. A. Neuroinflammatory astrocyte subtypes in the mouse brain. Nat Neurosci 24, 1475- 1487 (2021).

47. Marsh, S. E., et al. Dissection of artifactual and confounding glial signatures by single-cell sequencing of mouse and human brain. Nat Neurosci 25, 306–316 (2022).

48. Rotshenker, S. The role of Galectin-3/MAC-2 in the activation of the innate-immune function of phagocytosis in microglia in injury and disease. J Mol Neurosci 39, 99–103 (2009).

49. Marzan, D. E., et al. Activated microglia drive demyelination via CSF1R signaling. Glia 69, 1583–1604 (2021).

50. Zirngibl, M., Assinck, P., Sizov, A., Caprariello, A. V. & Plemel, J. R. Oligodendrocyte death and myelin loss in the cuprizone model: an updated overview of the intrinsic and extrinsic causes of cuprizone demyelination. Mol Neurodegener 17, 34 (2022).

51. Green, K. N., Crapser, J. D. & Hohsfield, L. A. To Kill a Microglia: A Case for CSF1R Inhibitors. Trends Immunol 41, 771–784 (2020).

52. Nikic, I., et al. A reversible form of axon damage in experimental autoimmune encephalomyelitis and multiple sclerosis. Nat Med 17, 495–499 (2011).

53. Yong, Y., Hunter-Chang, S., Stepanova, E. & Deppmann, C. Axonal spheroids in neurodegeneration. Mol Cell Neurosci 117, 103679 (2021).

54. Conforti, L., Gilley, J. & Coleman, M. P. Wallerian degeneration: an emerging axon death pathway linking injury and disease. Nat Rev Neurosci 15, 394–409 (2014).

55. Jung, J., et al. Actin polymerization is essential for myelin sheath fragmentation during Wallerian degeneration. J Neurosci 31, 2009–2015 (2011).

56. Catenaccio, A., et al. Molecular analysis of axonal-intrinsic and glial-associated co-regulation of axon degeneration. Cell Death Dis 8, e3166 (2017).

57. Vaquie, A., et al. Injured Axons Instruct Schwann Cells to Build Constricting Actin Spheres to Accelerate Axonal Disintegration. Cell Rep 27, 3152–3166 e3157 (2019).

58. Locatelli, G., et al. Mature oligodendrocytes actively increase in vivo cytoskeletal plasticity following CNS damage. J Neuroinflammation 12, 62 (2015).

59. Nawaz, S., et al. Actin filament turnover drives leading edge growth during myelin sheath formation in the central nervous system. Dev Cell 34, 139–151 (2015).

60. Sebbagh, M., Hamelin, J., Bertoglio, J., Solary, E. & Breard, J. Direct cleavage of ROCK II by granzyme B induces target cell membrane blebbing in a caspase-independent manner. J Exp Med 201, 465–471 (2005).

61. Ritter, A. T., et al. ESCRT-mediated membrane repair protects tumor-derived cells against T cell attack. Science 376, 377–382 (2022).

62. Costa, A. R., et al. The membrane periodic skeleton is an actomyosin network that regulates axonal diameter and conduction. Elife 9 (2020).

63. Berger, S. L., et al. Localized Myosin II Activity Regulates Assembly and Plasticity of the Axon Initial Segment. Neuron 97, 555–570 e556 (2018).

64. Bucur, O., et al. Nanoscale imaging of clinical specimens using conventional and rapid-expansion pathology. Nat Protoc 15, 1649–1672 (2020).

65. Thiery, J., et al. Perforin pores in the endosomal membrane trigger the release of endocytosed granzyme B into the cytosol of target cells. Nat Immunol 12, 770–777 (2011).

66. Edgar, J. M., et al. Oligodendroglial modulation of fast axonal transport in a mouse model of hereditary spastic paraplegia. J Cell Biol 166, 121–131 (2004).

67. Milde, S., Adalbert, R., Elaman, M. H. & Coleman, M. P. Axonal transport declines with age in two distinct phases separated by a period of relative stability. Neurobiol Aging 36, 971–981 (2015).

68. Loreto, A., et al. Mitochondrial impairment activates the Wallerian pathway through depletion of NMNAT2 leading to SARM1-dependent axon degeneration. Neurobiol Dis 134, 104678 (2020).

69. Simon, D. J., et al. Axon Degeneration Gated by Retrograde Activation of Somatic Pro-apoptotic Signaling. Cell 164, 1031–1045 (2016).

70. Schirmer, L., et al. Neuronal vulnerability and multilineage diversity in multiple sclerosis. Nature 573, 75–82 (2019).

71. Kenigsbuch, M., et al. A shared disease-associated oligodendrocyte signature among multiple CNS pathologies. Nat Neurosci 25, 876–886 (2022).

72. Pandey, S., et al. Disease-associated oligodendrocyte responses across neurodegenerative diseases. Cell Rep 40, 111189 (2022).

73. Bodini, B., et al. Dynamic Imaging of Individual Remyelination Profiles in Multiple Sclerosis. Ann Neurol 79, 726–738 (2016).

74. Mei, F., et al. Accelerated remyelination during inflammatory demyelination prevents axonal loss and improves functional recovery. Elife 5 (2016).

75. Ricigliano, V. A. G., et al. Spontaneous remyelination in lesions protects the integrity of surrounding tissues over time in multiple sclerosis. Eur J Neurol 29, 1719–1729 (2022).

76. Schäffner, E., et al. Myelin insulation as a risk factor for axonal degeneration in autoimmune demyelinating disease. Nat Neurosci (2023).

77. Nocera, G., et al. Repair oligodendrocytes demyelinating and disintegrating damaged axons after injury. bioRxiv, 2023.2005.2018.541273 (2023).

78. Bai, X., et al. In the mouse cortex, oligodendrocytes regain a plastic capacity, transforming into astrocytes after acute injury. Dev Cell (2023).

79. Jarjour, A. A., et al. The formation of paranodal spirals at the ends of CNS myelin sheaths requires the planar polarity protein Vangl2. Glia 68, 1840–1858 (2020).

80. Brown, T. L. & Macklin, W. B. The Actin Cytoskeleton in Myelinating Cells. Neurochem Res 45, 684–693 (2020).

81. Howell, O. W., et al. Activated microglia mediate axoglial disruption that contributes to axonal injury in multiple sclerosis. J Neuropathol Exp Neurol 69, 1017–1033 (2010).

82. Koehnle, T. J. & Brown, A. Slow axonal transport of neurofilament protein in cultured neurons. J Cell Biol 144, 447–458 (1999).

83. Unsain, N., et al. Remodeling of the Actin/Spectrin Membrane-associated Periodic Skeleton, Growth Cone Collapse and F-Actin Decrease during Axonal Degeneration. Sci Rep 8, 3007 (2018).

84. Datar, A., et al. The Roles of Microtubules and Membrane Tension in Axonal Beading, Retraction, and Atrophy. Biophys J 117, 880–891 (2019).

85. Griffiths, I., et al. Axonal swellings and degeneration in mice lacking the major proteolipid of myelin. Science 280, 1610–1613 (1998).

86. Babiychuk, E. B., Monastyrskaya, K., Potez, S. & Draeger, A. Blebbing confers resistance against cell lysis. Cell Death Differ 18, 80–89 (2011).

87. Klein, D. & Martini, R. Myelin and macrophages in the PNS: An intimate relationship in trauma and disease. Brain Res 1641, 130–138 (2016).

88. Malinarich, F., et al. High mitochondrial respiration and glycolytic capacity represent a metabolic phenotype of human tolerogenic dendritic cells. J Immunol 194, 5174–5186 (2015).

89. Gonzalez, M. A., et al. Phagocytosis increases an oxidative metabolic and immune suppressive signature in tumor macrophages. J Exp Med 220 (2023).

90. Safaiyan, S., et al. Age-related myelin degradation burdens the clearance function of microglia during aging. Nat Neurosci 19, 995–998 (2016).

91. Ip, C. W., et al. Origin of CD11b+ macrophage-like cells in the CNS of PLP- overexpressing mice: low influx of haematogenous macrophages and unchanged blood-brain-barrier in the optic nerve. Mol Cell Neurosci 38, 489–494 (2008).

92. Mombaerts, P., et al. RAG-1-deficient mice have no mature B and T lymphocytes. Cell 68, 869–877 (1992).

93. Milde, S., Fox, A. N., Freeman, M. R. & Coleman, M. P. Deletions within its subcellular targeting domain enhance the axon protective capacity of Nmnat2 in vivo. Sci Rep 3, 2567 (2013).

94. Zheng, G. X., et al. Massively parallel digital transcriptional profiling of single cells. Nat Commun 8, 14049 (2017).

95. Satija, R., Farrell, J. A., Gennert, D., Schier, A. F. & Regev, A. Spatial reconstruction of single-cell gene expression data. Nat Biotechnol 33, 495–502 (2015).

96. Zhou, Y., et al. Metascape provides a biologist-oriented resource for the analysis of systems-level datasets. Nat Commun 10, 1523 (2019).

97. Wang, F., et al. RNAscope: a novel in situ RNA analysis platform for formalin-fixed, paraffin-embedded tissues. J Mol Diagn 14, 22–29 (2012).

98. Emery, B. & Dugas, J. C. Purification of oligodendrocyte lineage cells from mouse cortices by immunopanning. Cold Spring Harb Protoc 2013, 854–868 (2013).

99. Andrews, S., Gilley, J. & Coleman, M. P. Difference Tracker: ImageJ plugins for fully automated analysis of multiple axonal transport parameters. J Neurosci Methods 193, 281–287 (2010).

100. Tonges, L., et al. Rho kinase inhibition modulates microglia activation and improves survival in a model of amyotrophic lateral sclerosis. Glia 62, 217–232 (2014).

101. Tatenhorst, L., et al. Fasudil attenuates aggregation of alpha-synuclein in models of Parkinson’s disease. Acta Neuropathol Commun 4, 39 (2016).

102. Faul, F., Erdfelder, E., Lang, A. G. & Buchner, A. G*Power 3: a flexible statistical power analysis program for the social, behavioral, and biomedical sciences. Behav Res Methods 39, 175–191 (2007).

