## Supplementary Figures and Legends for "Genetically perturbed myelin as a risk factor for neuroinflammation-driven axon degeneration"

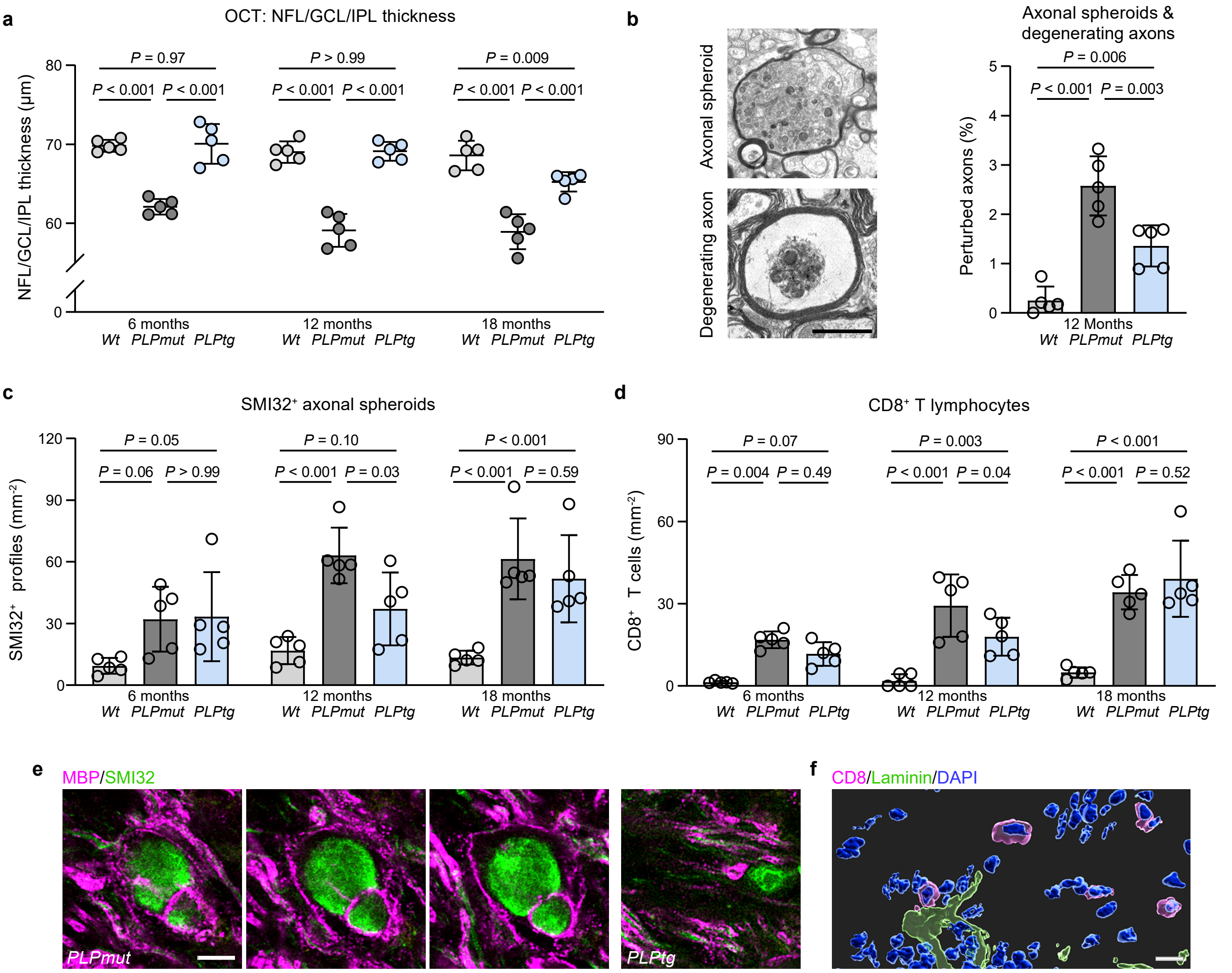


**Supplementary Fig. 1 I Similarities and differences of T cell-driven axonal damage in mice with distinct myelin defects. a** OCT analysis of the innermost retinal composite layer (NFL/GCL/IPL) in peripapillary circle scans of *Wt*, *PLPmut*, and *PLPtg* mice at different ages. Inner retinal thinning is much more pronounced in *PLPmut* than in *PLPtg* mice ($n$ = 5 mice per group, two-way ANOVA with Tukey's multiple comparisons test, $F$ (2, 36) = 133.4, $P$ < 0.001. **b** Electron microscopy-based quantification of axonal spheroids and degenerating axons in optic nerves from 12-month-old *Wt*, *PLPmut*, and *PLPtg* mice. Axonal damage is more pronounced in *PLPmut* than in *PLPtg* mice ($n$ = 5 mice per group, one-way ANOVA with Tukey's multiple comparisons test, $F$ (2, 12) = 32.91, $P$ < 0.001. **c** Quantification of axonal spheroids in optic nerves from *Wt*, *PLPmut*, and *PLPtg* mice at different ages (each circle represents the mean value of one mouse). $SMI32^+$ spheroids progressively accumulate in both *PLPmut* and *PLPtg* mice ($n$ = 5 mice per group, two-way ANOVA with Tukey's multiple comparisons test, $F$ (2, 36) = 25.66, $P$ < 0.001). **d** Quantification of $CD8^+$ T cells in optic nerves from *Wt*, *PLPmut*, and *PLPtg* mice at different ages (each circle represents the mean value of one mouse). $CD8^+$ T cells progressively accumulate in both *PLPmut* and *PLPtg* mice ($n$ = 5 mice per group, two-way ANOVA with Tukey's multiple comparisons test, $F$ (2, 36) = 50.4, $P$ < 0.001). **e** Immunofluorescence detection of SMI32 and MBP in the optic nerves of 12-month-old *PLPmut* and *PLPtg* mice by confocal microscopy. Individual optical slices show the presence of myelin around axonal spheroids in *PLPmut* mice. The smaller spheroids in *PLPtg* mice are often demyelinated. Scale bar, 5 µm. **f** IMARIS Z-stack surface rendering of $CD8^+$ T cells and $laminin^+$ blood vessels. Almost all $CD8^+$ T cells in optic nerves of myelin mutants are parenchymal. Scale bar, 10 µm. Data are presented as the mean ± s.d.

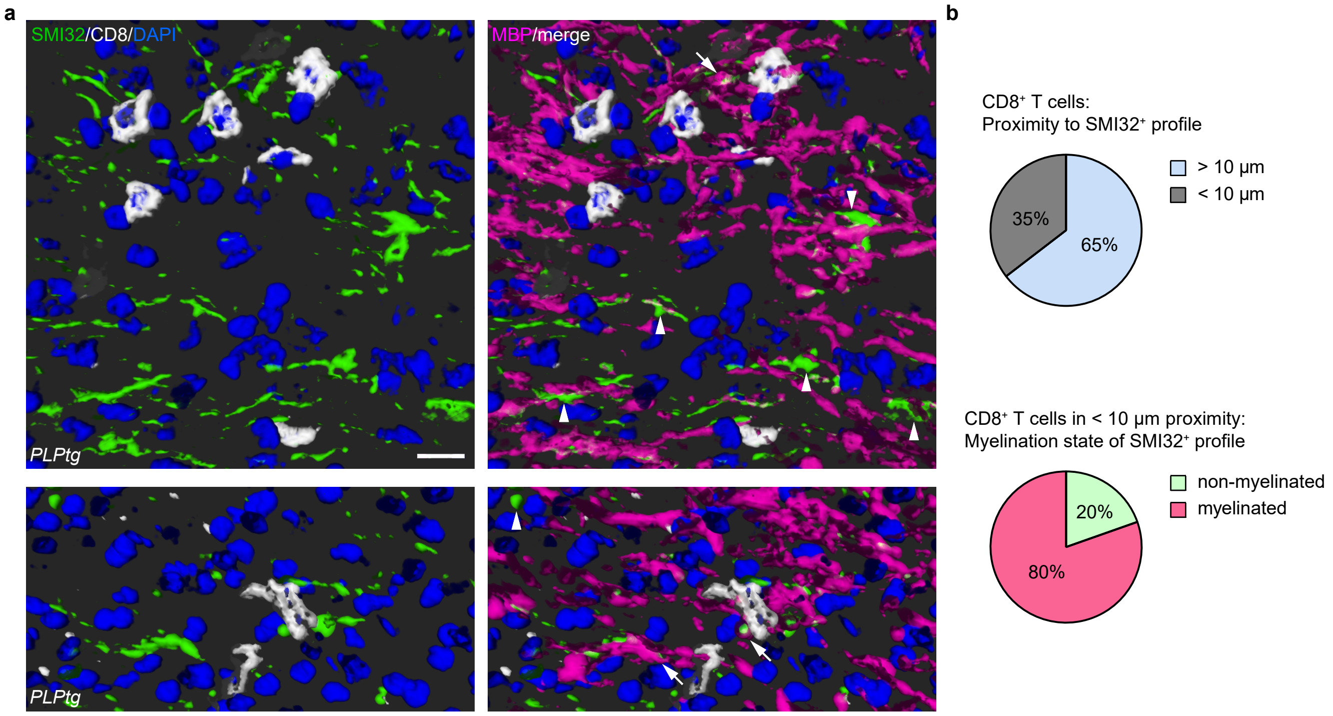


**Supplementary Fig. 2 I CD8⁺ T cells preferentially associate with damaged axons still ensheathed by perturbed myelin.**
**a** Immunofluorescence detection and IMARIS Z-stack surface rendering of CD8 in combination with SMI32 and MBP in optic nerves from 12-month-old *PLPtg* mice. Arrowheads indicate demyelinated SMI32⁺ fibers and arrows indicate SMI32⁺ axonal spheroids ensheathed by MBP⁺ myelin segments. Scale bar, 10 µm. **b** Quantification of CD8⁺ T cells in proximity (< 10 µm) to SMI32⁺ profiles (top) and myelination state of the respective SMI32⁺ profiles (bottom). CD8⁺ T cells preferentially accumulate in proximity to damaged fibers that are myelinated ($n$ = 144 CD8⁺ T cells from 5 *PLPtg* mice).

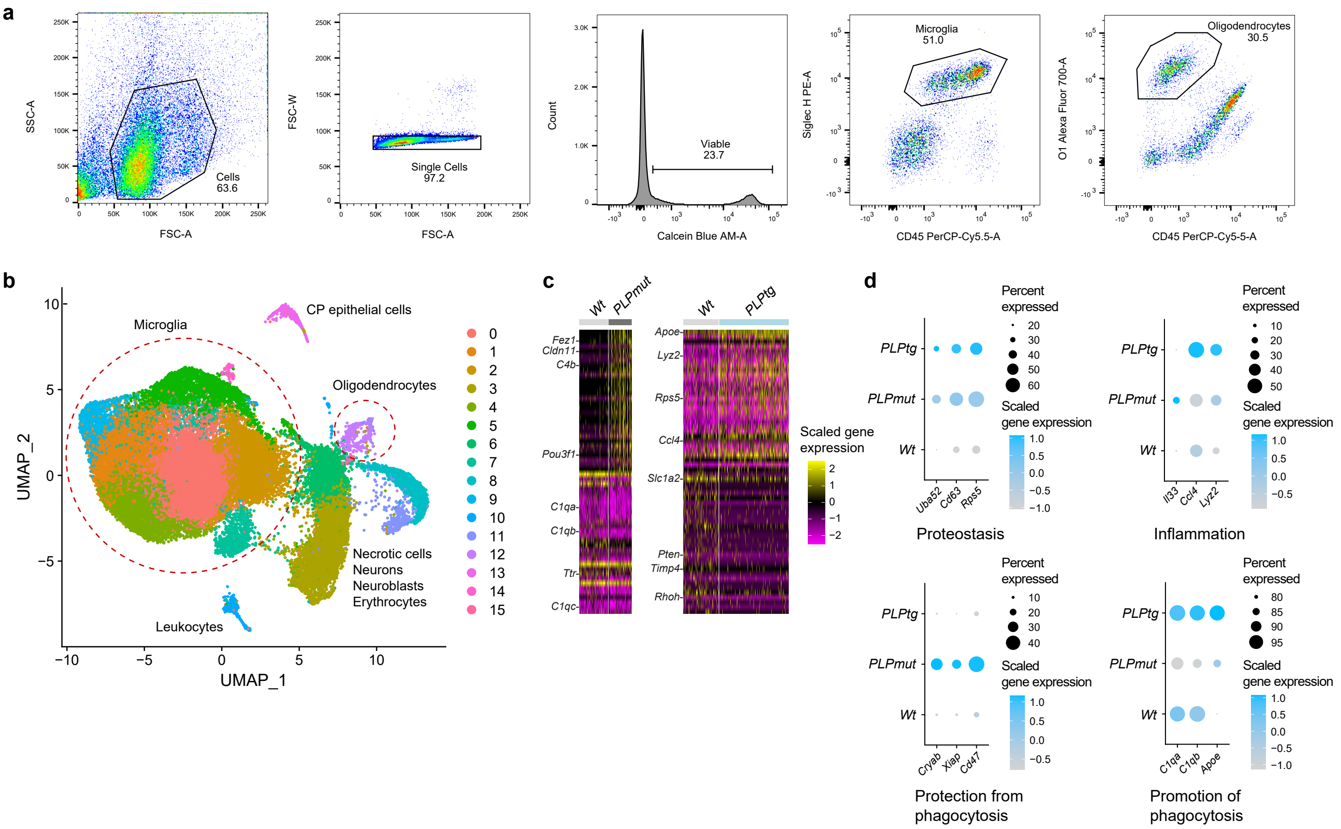


**Supplementary Fig. 3 I scRNA-seq of oligodendrocytes in *Wt*, *PLPmut*, and *PLPtg* mice reveals common and distinct disease-associated states. a** Representative FACS plots for gating of single, viable CD45$^{low}$Siglec-H$^{+}$ microglia and CD45$^{-}$O1$^{+}$ oligodendrocytes. **b** Combined UMAP visualization of 35,590 cells freshly sorted from adult (10-month-old) *Wt*, *PLPmut*, and *PLPtg* (*n* = 3 mice per group) mouse brains and analyzed by scRNA-seq. Microglia and oligodendrocyte clusters were identified based on marker gene expression, subsetted, and further analyzed as shown in Fig. 3a. **c** Heatmaps of top 30 differentially expressed genes comparing oligodendrocytes isolated from *Wt* and *PLPmut* (left) or *Wt* and *PLPtg* (right) brains as identified in Fig. 3a. The color scale is based on a z-score distribution from -2 (purple) to 2 (yellow). **d** Dot plot expression visualization of selected genes implicated in proteostasis (GO terms: cytoplasmic translation, lysosome); inflammation (GO terms: microglial activation involved in immune response, complement binding, lysozyme activity), protection from phagocytosis (GO terms: negative regulation of apoptotic process, negative regulation of Fc-gamma receptor signaling pathway involved in phagocytosis), or promotion of phagocytosis (GO terms: microglial cell activation, complement activation, cholesterol homeostasis) for oligodendrocytes as annotated in panel b. The color scales are based on z-score distributions from -1 or -0.5 (lightgrey) to 1 (lightblue). Complete lists of differentially expressed genes can be found in Supplementary Table 1.

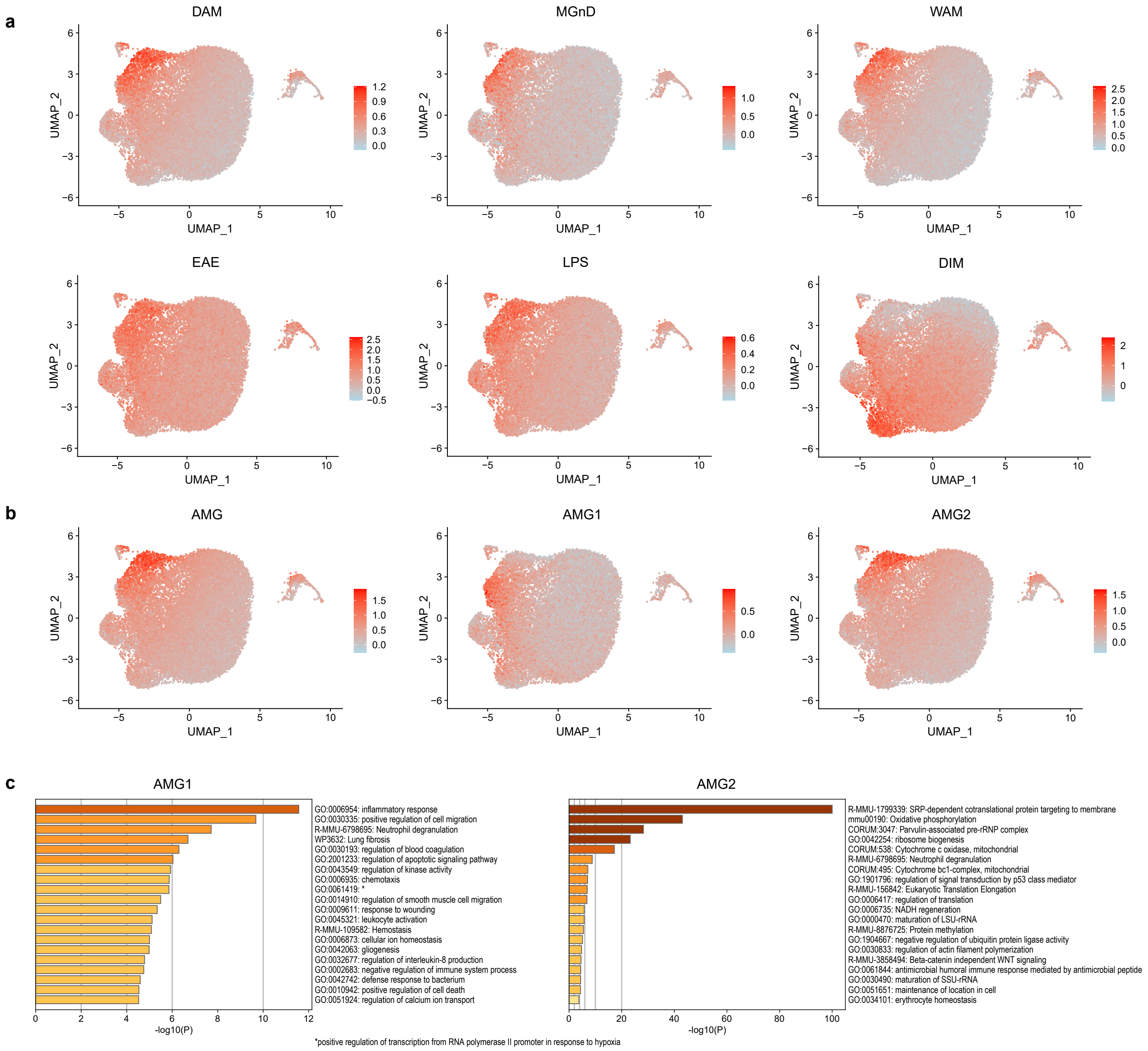


**Supplementary Fig. 4 I scRNA-seq reveals signatures of activated microglia in *PLPmut* and *PLPtg* mice. a** UMAP visualizations of MG and ODC (as annotated in Fig. 3a or Supplementary Fig. 3b) showing the expression of selected module scores. Transcript levels are color-coded: lightblue, not expressed; red, expressed. Module scores are based on top 300 upregulated genes of known microglia states as previously reported. **b** Module scores are based on identified upregulated genes shared between both AMG clusters, or enriched in either AMG1 or AMG2, respectively. **c** Metascape gene-set enrichment analysis of AMG1 or AMG2 module scores. Top 20 enriched terms are shown. Complete lists of module score genes can be found in Supplementary Table 2.

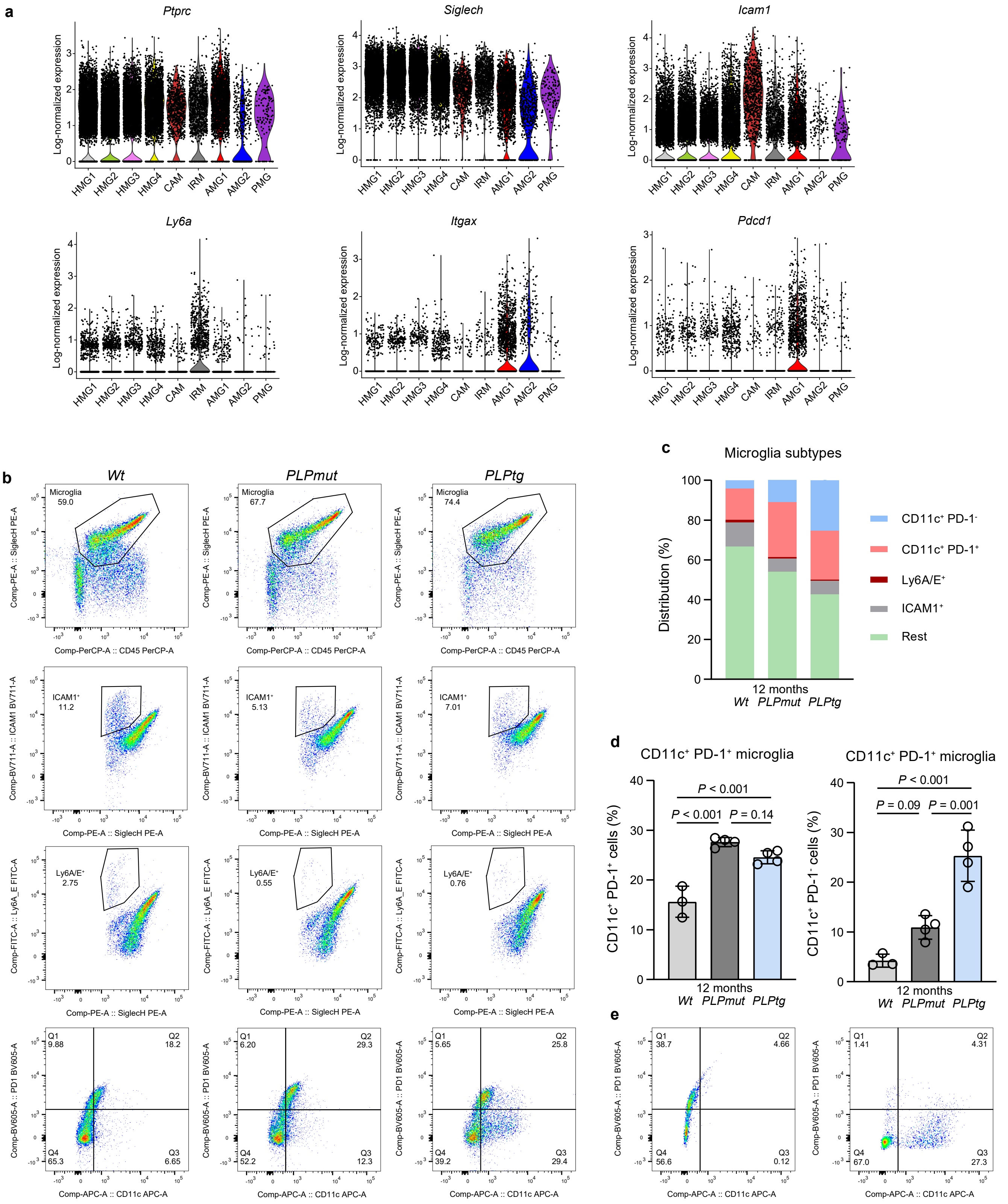

**Supplementary Fig. 5 I Flow cytometry confirms heterogeneity and distinct reactions of microglia in *PLPmut* and *PLPtg* mice.** **a** Violin plots of MG and ODC (as annotated in Fig. 3a) showing the expression of selected marker genes. Transcript levels are color-coded: lightblue, not expressed; red, expressed. **b** Representative flow cytometry plots for gating and analysis of single, viable CD45$^{low}$Siglec-H$^{+}$ microglia. Siglec-H$^{+}$ microglia subtypes comprise ICAM1$^{+}$ (CAM), Ly6A/E$^{+}$ (IRM), CD11c$^{+}$PD-1$^{+}$ (AMG1), and CD11c$^{+}$PD-1$^{-}$ (AMG2) cells. **c** Distribution of distinct microglia subtypes in brains from 12-month-old *Wt*, *PLPmut*, and *PLPtg* mice as identified in panel b. **d** Flow cytometry-based quantification of CD11c$^{+}$PD-1$^{+}$ (AMG1, left) and CD11c$^{+}$PD-1$^{-}$ (AMG2, right) populations among microglia. AMG1 is more frequent in both myelin mutants while AMG2 is significantly more frequent only in *PLPtg* mice ($n$ = 3-4 mice per group, one-way ANOVA with Tukey's multiple comparisons test, Left: $F$ (2, 8) = 38.10, $P$ < 0.001, Right: $F$ (2, 8) = 33.46, $P$ < 0.001). **e** Fluorescence minus one controls for CD11c (left) and PD-1 (right).

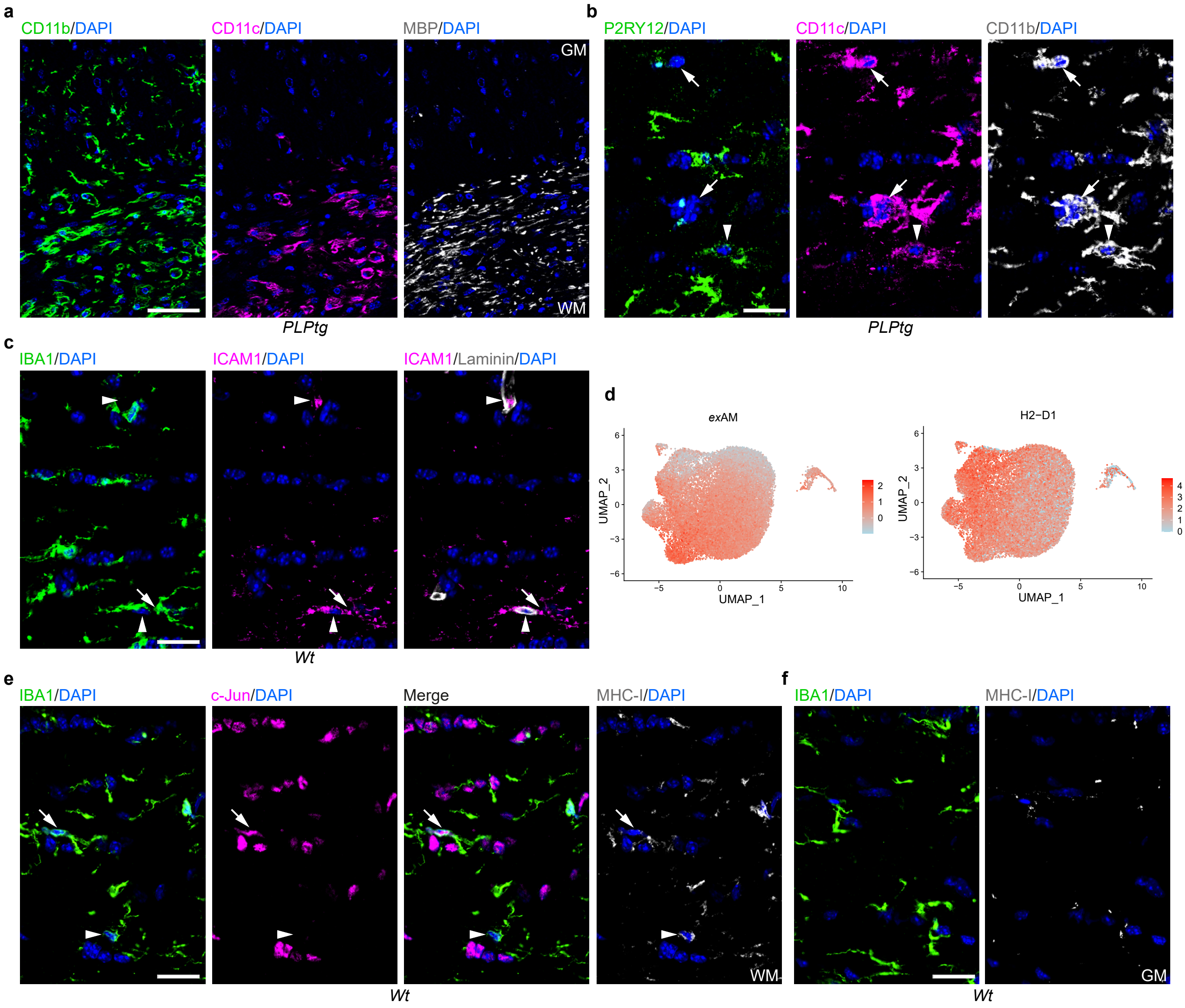


**Supplementary Fig. 6 I Microglial diversity is driven by localization and disease context. a** Representative immunofluorescence detection of CD11b in combination with CD11c and MBP in corpus callosum and lower cortex from 12-month-old *PLPtg* mice. Activated microglia (CD11b$^+$/CD11c$^+$) are detectable in white matter (WM) with ongoing demyelination but not in gray matter (GM). Scale bar, 50 µm. **b** Representative immunofluorescence detection of P2RY12 in combination with CD11c and CD11b in optic nerves from 12-month-old *PLPtg* mice. The arrowhead indicates a CD11b$^+$CD11c$^+$P2RY12$^+$ (AMG1) cell and the arrows indicate CD11b$^+$CD11c$^+$P2RY12$^-$ (AMG2) cells. Scale bar, 20 µm. **c** Representative immunofluorescence detection of IBA1 in combination with ICAM1 and Laminin in optic nerves from 12-month-old *Wt* mice. The arrowheads indicate IBA1$^-$ICAM1$^+$Laminin$^+$ capillaries and the arrow indicates an IBA1$^+$ICAM1$^+$Laminin$^-$ capillary-associated microglial cell (CAM). Scale bar, 20 µm. **d** UMAP visualizations of MG and ODC (as annotated in Fig. 3a) showing the expression of the exAM module score[44] or *H2-D1*. Transcript levels are color-coded: lightblue, not expressed; red, expressed. **e** Representative immunofluorescence detection of IBA1 in combination with c-Jun and MHC-I in optic nerves from 12-month-old *Wt* mice. The arrowhead indicates an IBA1$^+$c-Jun$^-$MHC-I$^{low}$ (HMG4) cell and the arrow indicates an IBA1$^+$c-Jun$^-$MHC-I$^{low}$ (HMG3) in white matter (WM). **f** Cortical gray matter (GM) microglia in adult *Wt* mice are IBA1$^+$MHC-I$^-$ (HMG1, HMG2). Scale bars, 20 µm.

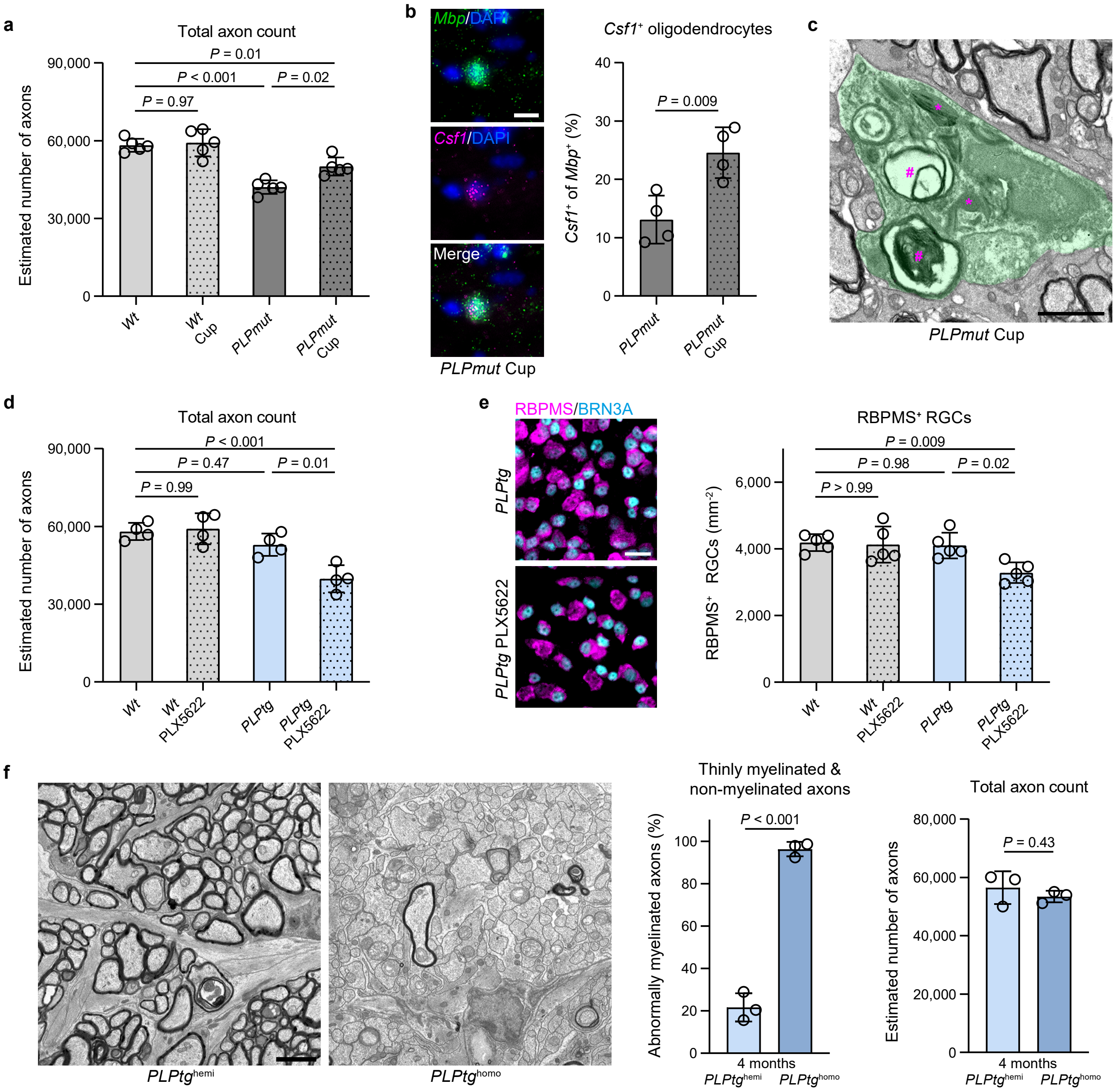


**Supplementary Fig. 7 I Modulating myelin phagocytosis inversely affects axon degeneration in distinct myelin mutants . a** Electron microscopy-based estimation of total axonal numbers in the optic nerves of *Wt* and *PLPmut* mice after control or cuprizone (Cup) diet (each circle represents the mean value of one mouse). Cuprizone significantly attenuates axon loss in *PLPmut* mice ($n$ = 5 mice per group, one-way ANOVA with Tukey's multiple comparisons test, $F$ (3, 16) = 24.38, $P$ < 0.001). **b** RNAscope *in situ* hybridization for *Mbp* and *Csf1* in the optic nerves of *PLPmut* mice after control or cuprizone (Cup) diet (each circle represents the mean value of one mouse). Cuprizone significantly increases the frequency of *Csf1*$^+$*Mbp*$^+$ oligodendrocytes in *PLPmut* mice. ($n$ = 5 mice per group, two-sided Student's t-test, $t$ = 3.834, d.f. = 6). Scale bar, 10 µm. **c** Electron microscopy demonstrates intracellular accumulation of myelin fragments (hashtags) and lysosomal storage material (asterisks) in optic nerves from *PLPmut* Cup mice. Scale bar, 2 µm. **d** Electron microscopy-based estimation of total axonal numbers in the optic nerves of *Wt* and *PLPtg* mice after control or PLX5622 diet (each circle represents the mean value of one mouse). PLX5622 significantly increases axon loss in *PLPtg* mice ($n$ = 4 mice per group, one-way ANOVA with Tukey's multiple comparisons test, $F$ (3, 12) = 13.56, $P$ < 0.001). **e** Immunofluorescence detection and quantification of RBPMS$^+$BRN3A$^+$ RGCs in the retinae of *Wt* and *PLPtg* mice after control or PLX5622 diet. PLX5622 causes neuron loss in *PLPtg* mice ($n$ = 5 mice per group, one-way ANOVA with Tukey's multiple comparisons test, $F$ (3, 16) = 6.156, $P$ = 0.006). Scale bar, 20 µm. **f** Representative electron micrographs of optic nerve cross-sections from 4-month-old hemizygous (left) or homozygous (right) *PLPtg* mice and quantification of thinly myelinated (g-ratio ≥ 0.85) and non-myelinated axons or total axonal numbers ($n$ = 3 mice per group, two-sided Student's t-test, myelination: $t$ = 17.16, d.f. = 4, axon count: $t$ = 0.8863, d.f. = 4). Scale bar, 10 µm. Data are presented as the mean ± s.d.

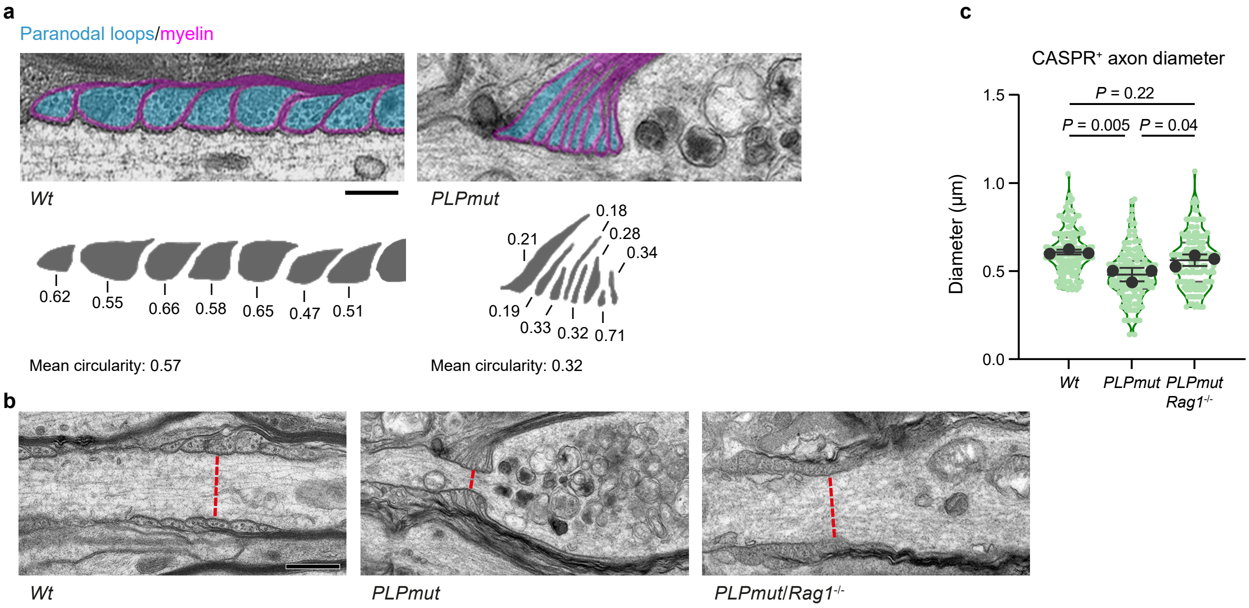


**Supplementary Fig. 8 I Decreased circularity of paranodal loops adjacent to axonal spheroids and reduced paranodal diameter in myelin mutant mice. a** Representative electron micrographs of paranodal loops in optic nerve longitudinal sections from 9-month-old *Wt* and *PLPmut* mice (top). Oligodendroglial cytoplasmic paranodal loops are pseudocolored in cyan and myelin membranes in magenta. Individual and mean paranodal loop cytoplasmic shapes and their circularity (form factor) are decreased in proximity to axonal spheroids in *PLPmut* mice (bottom). Quantifications are provided in Fig. 6d. Scale bar, 0.25 µm. **b** Representative electron micrographs of paranodal domains in optic nerve longitudinal sections from 9-month-old *Wt*, *PLPmut*, and *PLPmut*/*Rag1*$^{-/-}$ mice. Myelin compaction is abnormal in both *PLPmut* and *PLPmut*/*Rag1*$^{-/-}$ mice but minimum paranodal axon diameters (red dahed lines) are only decreased in *PLPmut* mice. Quantifications are provided in Fig. 6e. Scale bar, 0.5 µm. **c** Immunofluorescence-based measurement of CASPR$^{+}$ paranodal axon diameter in 12-month-old *Wt*, *PLPmut*, and *PLPmut*/*Rag1*$^{-/-}$ mice (green circles represent individual paranodes and black circles represent the mean of one mouse). Paranodal axon diameters are decreased in *PLPmut* but not *PLPmut*/*Rag1*$^{-/-}$ mice ($n$ = 50 paranodes per mouse and 3 mice per group, one-way ANOVA with Tukey's multiple comparisons test, $F$ (2, 6) = 14.08, $P$ = 0.005). Data are presented as the mean ± s.d.

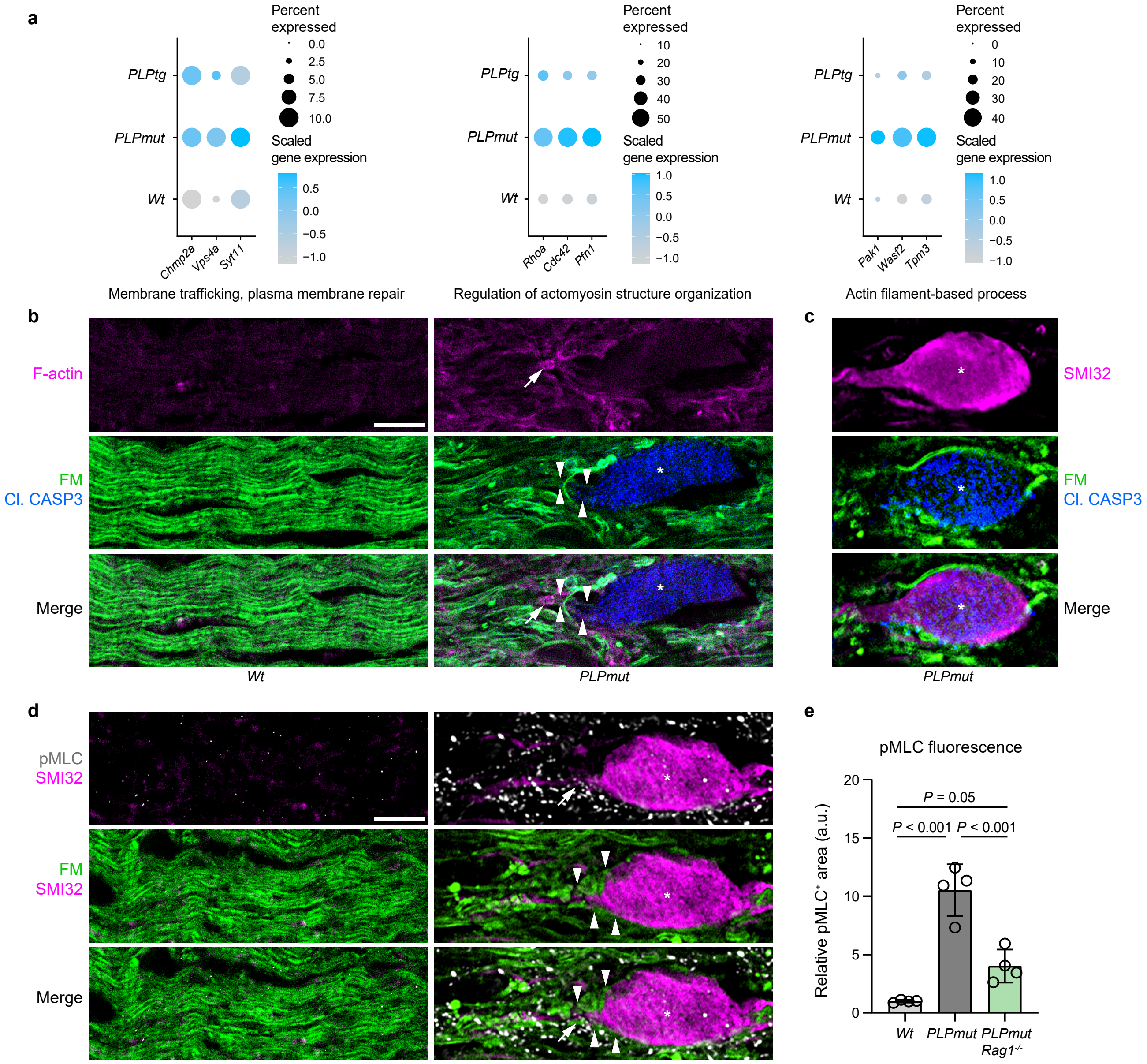


**Supplementary Fig. 9 I Axonal spheroid formation in myelin mutant mice is associated with altered cytoskeletal plasticity.**
**a** Dot plot expression visualization of selected genes implicated in GO terms membrane trafficking, plasma membrane repair, regulation of actomyosin structure organization, or actin filament-based process for oligodendrocytes as annotated in Figure 3a. The color scales are based on z-score distributions from -1 (lightgrey) to 1 (lightblue). Complete lists of differentially expressed genes can be found in Supplementary Table 1. **b** Representative immunofluorescence detection of F-actin (phalloidin) reactivity in combination with Fluoromyelin (FM) and cleaved caspase 3 in optic nerves from 12-month-old *Wt* (left) and *PLPmut* (right) mice. The asterisk indicates a Cl. CASP3$^+$ axonal spheroid, arrowheads indicate its ensheathing FM$^+$ myelin segment, and the arrow indicates an outer (putative paranodal) aspect with strong F-actin reactivity. **c** Immunofluorescence confirms caspase 3 activation in myelinated SMI32$^+$ axonal spheroids (asterisk) of *PLPmut* mice (98.44 % of SMI32$^+$ profiles are Cl. CASP3$^+$). **d** Representative immunofluorescence detection of pMLC reactivity in combination with SMI32 and FM in the optic nerves from 12-month-old *Wt* and *PLPmut* mice. The asterisk indicates an SMI32$^+$ axonal spheroid, arrowheads indicate its ensheathing FM$^+$ myelin segment, and the arrow indicates an outer (putative paranodal) aspect with strong pMLC reactivity. Scale bars, 10 µm. **e** Quantification of pMLC fluorescence by thresholding analysis demonstrates an increased relative pMLC$^+$ area in 12-month-old *PLPmut* mice and a significant attenuation of pMLC fluorescence in *PLPmut*/*Rag1*$^{-/-}$ mice (each circle represents the mean value of 1 mouse, $n$ = 4 mice per group, One-way ANOVA with Tukey's multiple comparisons test, $F$ (2, 9) = 41.00, $P$ < 0.001). Data are presented as the mean ± s.d.

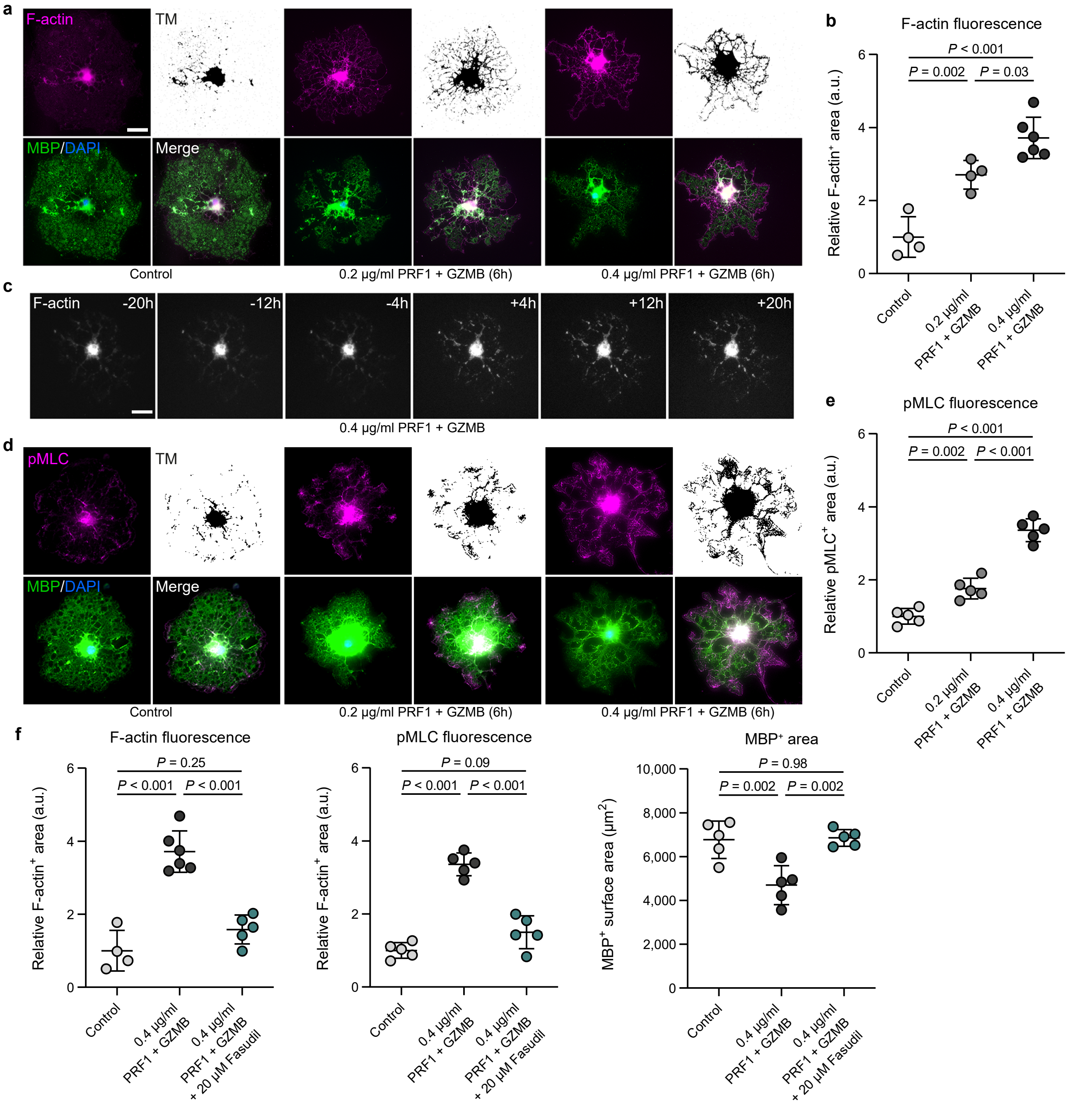


**Supplementary Fig. 10 I Cytotoxic effector molecules induce cytoskeletal plasticity and actomyosin constriction in myelinating oligodendrocytes. a** Representative immunofluorescence detection of F-actin (phalloidin) reactivity in combination with MBP in cultured ODC 6h after treatment with control medium (left) and medium containing 0.2 (middle) or 0.4 (right) µg/ml PRF1 + GZMB. TM: thresholding mask for analysis of F-actin fluorescence. Scale bar, 20 µm. **b** Quantification of F-actin fluorescence by thresholding analysis demonstrates a dose-dependent increased relative F-actin⁺ area after cytotoxic treatment (each circle represents the mean value of 5 ODC per well, $n$ = 3 independent experiments with 3 mice each, One-way ANOVA with Tukey's multiple comparisons test, $F$ (2, 11) = 32.42, $P$ < 0.001). **c** Representative live imaging of an ODC transfected with CellLight Actin-RFP at different time points before and after cytotoxic treatment. Scale bar, 20 µm. **d** Representative immunofluorescence detection of pMLC reactivity in combination with MBP in cultured ODC 6h after treatment with control medium (left) and medium containing 0.2 (middle) or 0.4 (right) µg/ml PRF1 + GZMB. TM: thresholding mask for analysis of pMLC fluorescence. **e** Quantification of F-actin fluorescence by thresholding analysis demonstrates a dose-dependent increased relative pMLC⁺ area after cytotoxic treatment ($n$ = 3 independent experiments with 3 mice each, One-way ANOVA with Tukey's multiple comparisons test, $F$ (2, 12) = 97.58, $P$ < 0.001). **f** Quantification of F-actin (left) or pMLC (middle) fluorescence and MBP (right) area by thresholding analysis after cytotoxic treatment (0.4 µg/ml) with or without Fasudil (20 µM). Fasudil inhibits the increase of F-actin and pMLC levels and the decrease in MBP⁺ surface area after PRF1 + GZMB ($n$ = 3 independent experiments with 3 mice each, One-way ANOVA with Tukey's multiple comparisons test, F-actin: $F$ (2, 12) = 40.40, $P$ < 0.001, pMLC: $F$ (2, 12) = 67.47, $P$ < 0.001, MBP: $F$ (2, 12) = 13.43, $P$ < 0.001). Data are presented as the mean ± s.d.
